# Lineage-specific change in craniofacial gene regulatory networks of syngnathid fishes revealed by integrating multiomics across fishes

**DOI:** 10.64898/2026.07.16.738491

**Authors:** Hope Healey, Clara Rehmann, Susan Bassham, William A. Cresko

## Abstract

Cranial neural crest (CNC) cells are essential developmental contributors to the remarkable diversity of vertebrate skull shapes, yet how underlying gene regulatory networks (GRNs) evolve to produce highly derived morphologies remains a challenging question. Syngnathid fishes (seahorses, pipefishes, pipehorses, and seadragons) are an opportune family of species in which to address this problem because of their unusual and extensive cranial diversity and their loss of craniofacial patterning genes, *fgf3* and *fgf4*. Here we investigated whether syngnathid craniofacial evolution experienced only a few localized network changes or required global rewiring of CNC GRNs. Using comparative single-cell RNA sequencing, ATAC-seq and whole genome alignments across Gulf pipefish, threespine stickleback, and zebrafish, we found that the core pharyngeal arch CNC gene network is notably conserved in syngnathids despite their derived morphology. However, we identified key local changes including expression of *fgf22* in percomorph CNC-derived pharyngeal arch cells that is not shared with more basally diverging zebrafish, as well as syngnathid-specific changes in conserved regulatory elements associated with the genes *ece1* and *spry2*. We propose that, while loss of *fgf3* expression causes severe craniofacial defects in zebrafish, pharyngeal CNC expression of *fgf22* in the percomorph fish lineage provided functional redundancy and relaxed constraint on *fgf3/4*, and that altered regulation of Fgf pathway modulators could contribute to craniofacial elaboration. Our findings support a model in which local GRN modifications, rather than widespread network rewiring, underlie the evolution of derived syngnathid craniofacial structures.

**Significance:** Understanding how developmental genetic changes drive the evolution of unique traits remains a long-standing challenge in biology. In the case of syngnathid fishes (pipefishes, seahorses, and seadragons), previous genomic studies identified candidate craniofacial gene losses which are proposed to relate to their elongate and derived heads, but the developmental impact of these losses is unknown. Through gene expression and comparative genomics analyses, we find that these fishes have distinct changes to craniofacial gene regulatory networks including gene content losses and gene expression gains and losses. Our study suggests that morphological adaptations may arise from multiple key changes within largely conserved developmental regulatory networks.

## Introduction

Despite tremendous craniofacial diversity across vertebrates, core regulatory genes are largely conserved in sequence, expression, and function (1). For instance, *dlx2* orthologs maintain expression in pharyngeal arch (PA) cells from basal vertebrates to mammals (2). These pharyngeal arch cells originate from the cranial neural crest (CNC) and contribute to most of the head and jaw. Craniofacial morphological changes can originate from gene regulatory network (GRN) re-wiring in CNC or interacting cells that impact cell survival, proliferation, and/or migration into pharyngeal arches or the frontonasal stream (3). Do these phenotypic changes require a small number of genetic changes, or more global rewiring? Some studies have identified changes in a single gene that is proposed to lead to craniofacial divergence. For example, SNPs in the gene *lbh* have been suggested to impact CNC migration in cichlids (4). However, in comparisons with larger evolutionary distance and similar craniofacial structures, numerous sequence changes in genes and regulatory elements influence CNC expression patterns (e.g.: ∼13% of Human CNCC enhancers have reduced or increased activity compared to Chimpanzee; (5). Therefore, the scope of CNC network changes in organisms with both high evolutionary divergence and derived heads is largely unknown.

Syngnathid fishes (seahorses, pipefishes, pipehorses, and seadragons) are an exciting system to investigate GRNs underlying derived craniofacial structures. Their heads are highly modified and include a tube-like snout which lacks teeth, and comprise of elongated ethmoid bones and a specialized hyoid that allows the head to rapidly pivot for prey capture (6, 7). These fishes have lost deeply conserved signaling genes (*fgf3, fgf4*), transcription factors (*eve1, tbx4*), and regulatory elements (*hoxa2b;* 8–10). The loss of *fgf3* is particularly surprising given that zebrafish *fgf3* mutants lack gill arches (PA 3-7 derivatives) and prematurely die (11, 12). Despite this loss of *fgf3*, syngnathids still possess gill arches. Therefore, CNC networks in syngnathids as compared to zebrafish must have differences in structure and/or composition beyond the described gene losses. We set out to discover these broader developmental genetic changes in syngnathid craniofacial development.

Given the elongated snouts and loss of *fgf3* of syngnathids, we asked the extent (local vs. global) to which craniofacial gene network re-wiring has occurred during the evolution of syngnathids. In addition to the extent, we investigated the mode of evolutionary changes. Gene networks can be modified through several evolutionary mechanisms including: changes in gene regulatory elements; gene loss, gain or co-option; and gene or genome duplication (13, 14). Gene duplicates, such as those originating from the teleost whole genome duplication event, can have differential fates, such as gene loss, subfunction partitioning, and acquisition of novel function any of which might lead to phenotypic evolution (15–17). Under the local re-wiring hypothesis, we propose that one or two genetic changes led to redundancy with *fgf3/fgf4* prior to their loss and these same or a few additional alterations promoted craniofacial elongation. In the global re-wiring hypothesis, we consider that extensive, global re-wiring in the CNC network reduced the importance of Fgf pathways in CNC cells and led to a derived head. The two mentioned hypotheses exist on opposite ends of a continuum of GRN modifications, with possibility for the true scenario to be within the spectrum from strict local or global hypotheses.

To detect changes in syngnathid craniofacial GRNs, and to determine whether these changes are more extensive than in other cell lineages, we harnessed several comparative “omics” approaches including single cell RNA sequencing (scRNAseq), whole genome alignments (WGA), and Assay for Transposase-Accessible Chromatin sequencing (ATAC-seq). We created sequencing libraries from a syngnathid (*Syngnathus scovelli*, Gulf pipefish) and another percomorph fish species (*Gasterosteus aculeatus*, threespine stickleback). We then integrated these data with published data from the more distantly related zebrafish (*Danio rerio*). We selected these teleosts to include a syngnathid (Gulf pipefish) and two fish with conventional snouts (threespine stickleback and zebrafish) while also accounting for phylogenetic relationships (zebrafish is a cypriniform and stickleback and pipefish are percomorphs). Although we detected in a select set of genes changes in CNC expression, presence in the genome, and/or regulatory elements, overall, we found that the core syngnathid PA CNC GRN is conserved. These results support that syngnathid craniofacial gene network and morphological evolution may have occurred as local changes in GRNs rather than global re-wiring, increasing the probability that causative genetic changes could ultimately be inferred with greater confidence.

## Results

### Successful integration of Gulf pipefish, stickleback and zebrafish single cell data sets

To compare syngnathid craniofacial gene expression to species with non-derived heads and intact *fgf3* and *4* genes, we completed a cross-species single cell analysis using Gulf pipefish, stickleback, and zebrafish at the pharyngeal arch stage. This is the stage during which loss of *fgf3* is detrimental to zebrafish head development (11). The integrated atlas contained 32,814 cells (5,469 cells per sample) that contributed to 47 cell clusters (Fig. 1). From cluster annotation with *de novo* marker genes determined using published zebrafish resources (Fig. S1; 18–21), we identified cells from all major cell type lineages. These included neural (such as CNC derived neurons), somite, mesenchyme, epidermis, periderm, pigment, immune, erythrocyte, pharyngeal pouch, and CNC pharyngeal arch cells. We evaluated species’ cell mixing as well as cell cluster separation using iLISI scores, a metric for cell proximity in UMAP space. Apart from erythrocyte clusters that are not pertinent for this analysis, species’ cells mixed well in UMAP space, suggesting a successfully integrated atlas (Fig. 2; 22). In addition, the cell clusters were well distinguished from one another, enabling analyses comparing these clusters (Fig. S2).

**Fig. 1.**
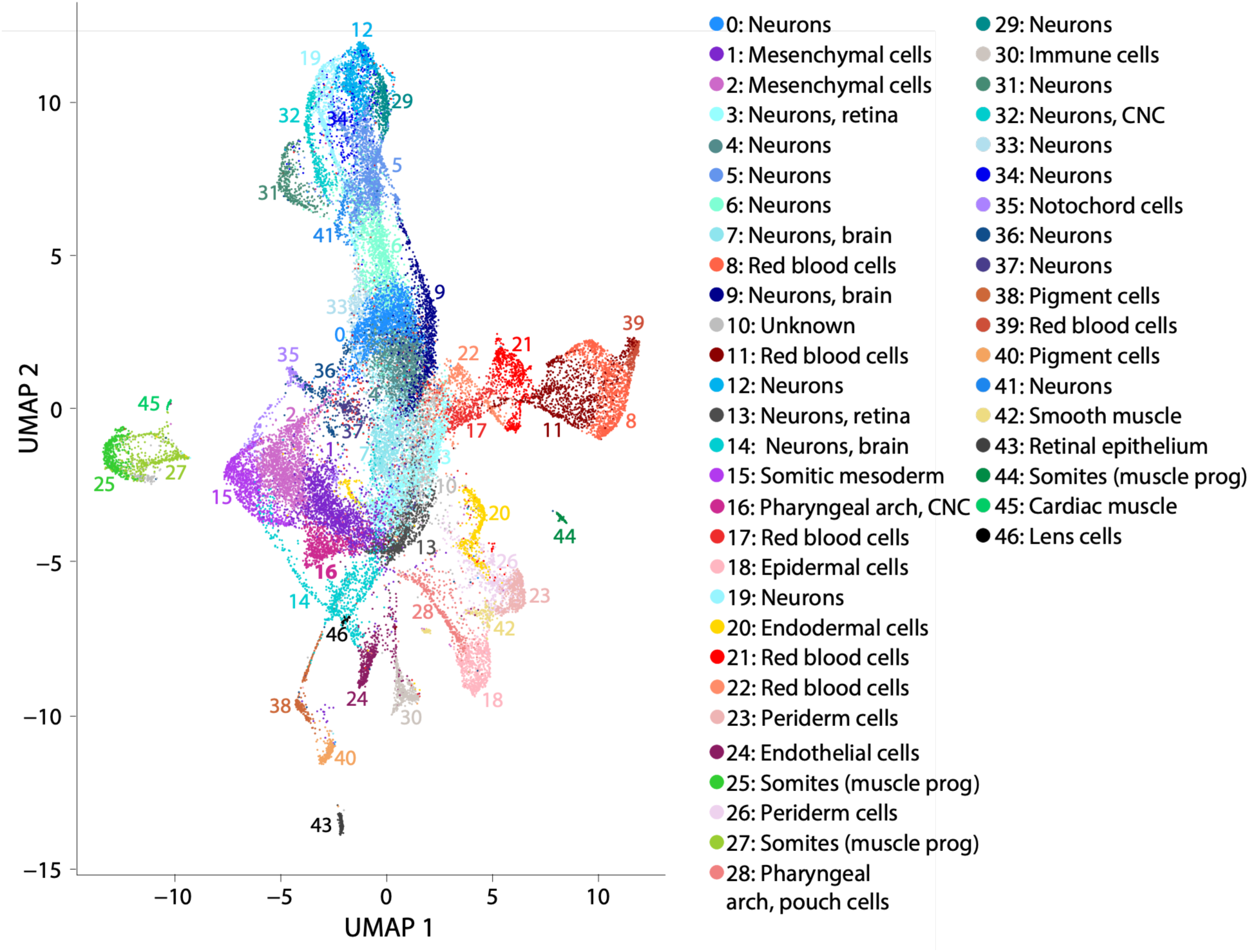
Integration of three scRNAseq atlases and identification of PA CNC cells. Cell types were identified from 46 cell clusters of the scRNAseq atlas (panel A), including CNC pharyngeal arch cells in cluster 16.

**Fig. 2.**
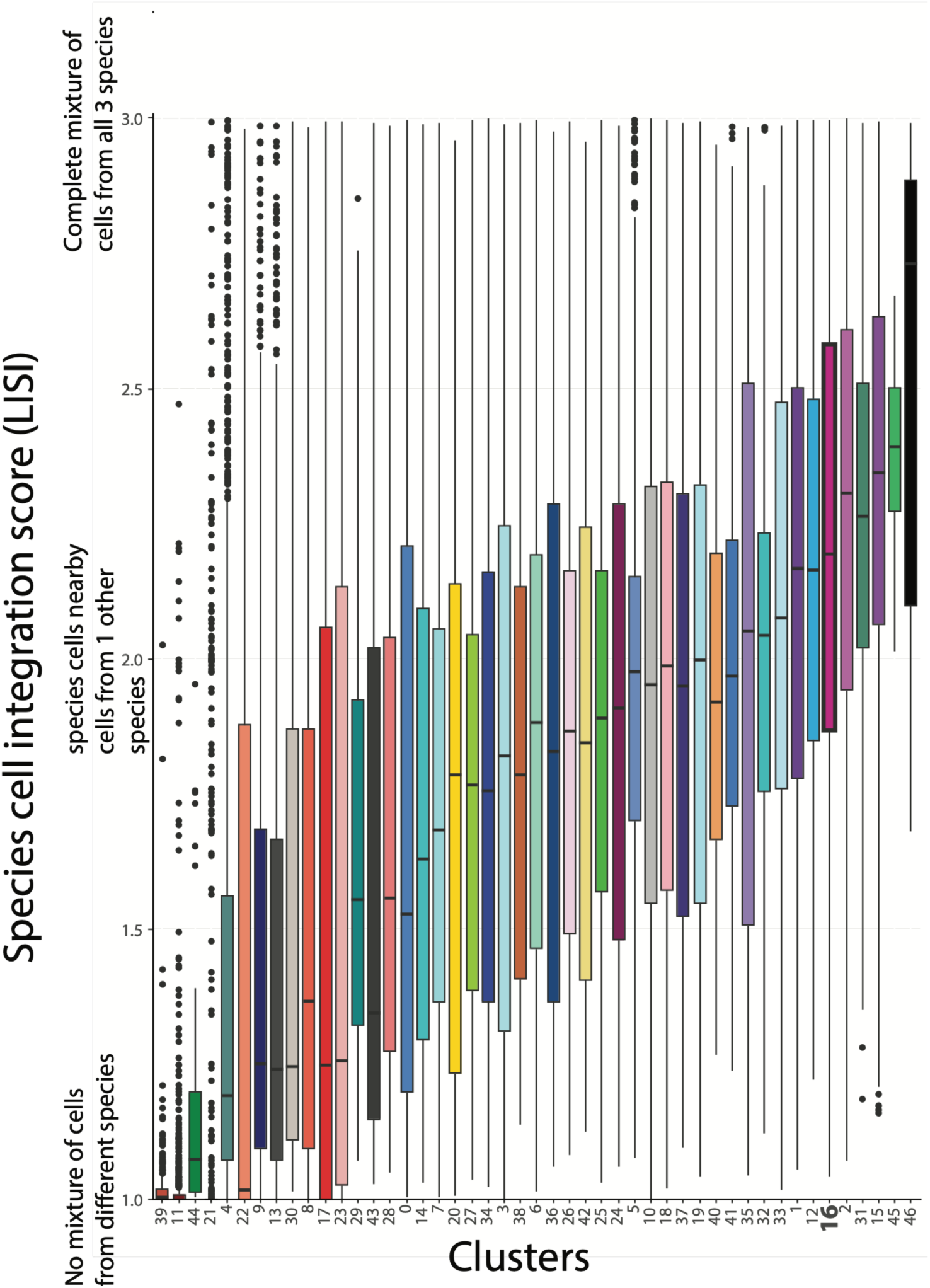
CNC PA cells from 3 species are well mixed. Integrated success was measured using the LISI metric and evaluated across all cell clusters. For an atlas with three species, an iLISI score of 3 indicates perfect integration and a score of 1 suggests failed integration.

### Cranial neural crest pharyngeal arch core gene regulatory network is conserved in Gulf pipefish

To test our hypothesis that syngnathid CNC PA cells have highly altered gene networks, we used metrics for global expression to compare Gulf pipefish CNC PA cells with the corresponding cells in stickleback and zebrafish. We first compared the correlation of cell type gene expression patterns across species and then asked whether the strength of association was reduced for pipefish in comparisons of the pharyngeal arch cell type. For this analysis, we identified cellular neighborhoods from each species’ data, and then completed a pairwise cellular neighborhood correlation analysis with 3,320 highly variable 1:1:1 orthologs (23). Across all 47 cell clusters, the PA CNC cluster was highly conserved (within the top six highly correlative clusters; Tables S2). To reduce bias from species differences in cell abundances, we binned clusters into major cell types. Once again, pharyngeal arch cells were one of the cell types with the highest correlation across all comparisons (Fig. 3A-C). The median correlation scores of PA cells across the three comparisons were congruent with the phylogeny: pipefish-stickleback (.87), stickleback-zebrafish (.85), and pipefish-zebrafish (.80; Fig. S3). Overall, these data support a high degree of similarity between stickleback and pipefish CNC PA cell gene expression patterns.

**Fig. 3.**
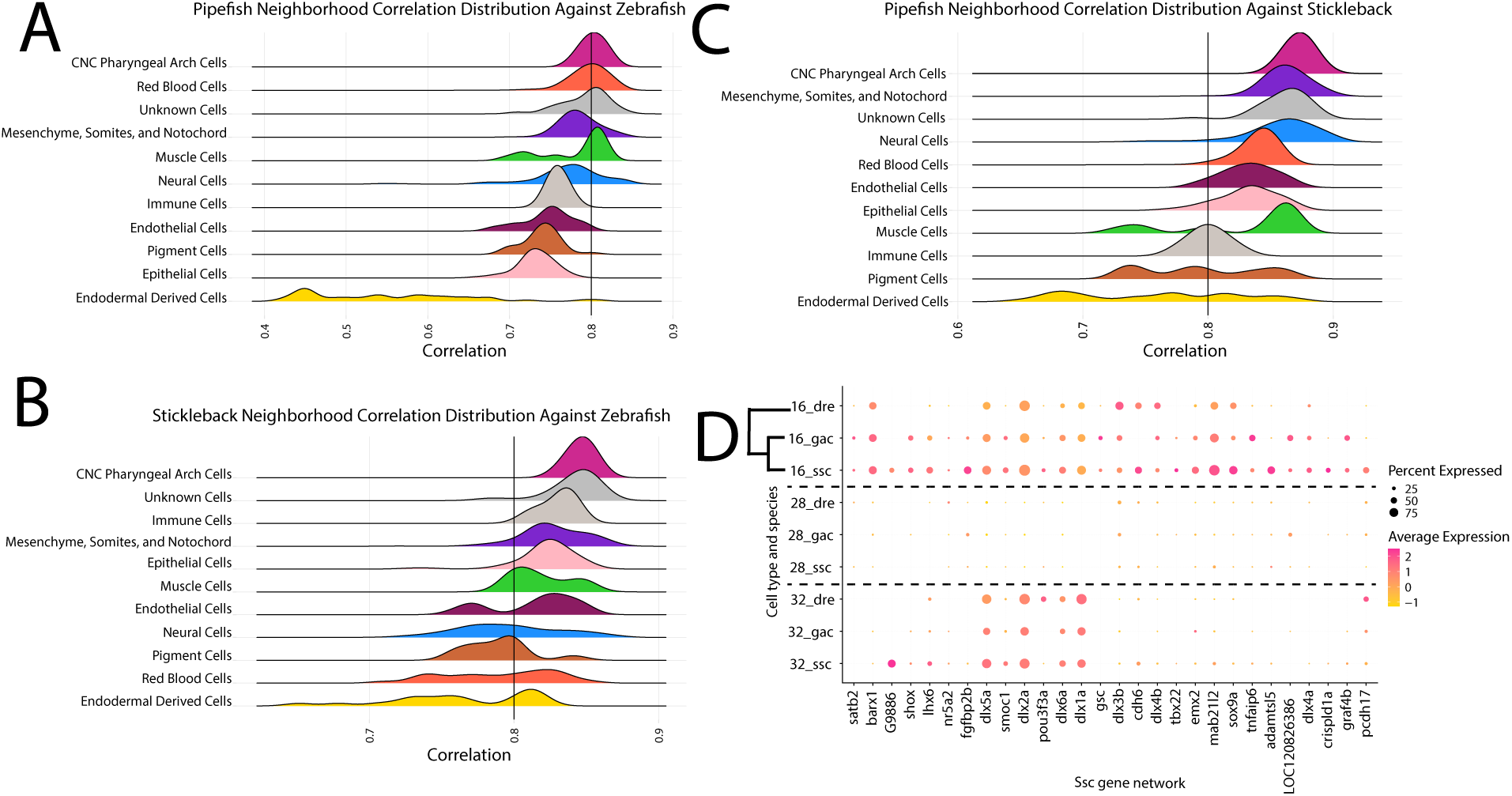
Pharyngeal arch cells express highly conserved core genetic program. In panels A-C, pairwise correlation analyses demonstrate that pharyngeal arch cells have maintained core gene expression profile in all species. Cell types are binned into categories based on common identities. A black line denotes a correlation of 80%. In panel D, the genes from the Gulf pipefish genetic network that correlate with pharyngeal arch cells are plotted. Cell types are separated by species (dre, zebrafish; gac, stickleback; ssc, Gulf pipefish). The size of the dot is proportional to the percentage of cells that express the gene, and the average expression is denoted via color. Cluster 16, cranial neural crest pharyngeal arch cells, cluster 28, pharyngeal pouch cells, and cluster 32, CNC neural derived cells, are shown on the plot.

Based on this preliminary evidence of gene network conservation, we investigated PA cell gene network composition using WGCNA (24). We built gene networks in the integrated atlas and individual species’ atlases. However, we only recovered a PA network in the pipefish, perhaps due to a greater proportion of CNC PA cells in the pipefish atlas. We compared expression of pipefish PA network genes across species (Fig. 3D) and observed a high level of similarity in presence/absence of these genes’ expression across species: *dlx* genes, *barx1, lhx6, mab21l2, sox9a*, and *emx2*. The exception to this pattern was *G9886,* which corresponds to an ortholog in stickleback (*LOC120826634*), but the stickleback gene was removed during Cell Ranger’s filtering steps because it is annotated as a ncRNA. Thus, the uniqueness of *G9886’s* expression in PA cells is unknown. Similar to results of the cross-species correlation analysis, this WGCNA finding suggests that the PA network detected in pipefish is largely conserved.

### Evolution of key developmental genes related to craniofacial networks

With no evidence for global shifts in Gulf pipefish CNC PA expression networks, we next used comparative ‘omics techniques to evaluate the extent of gene loss, gene gain, and conserved element change in syngnathids that could impact craniofacial development. We, first, identified genes that were not detected in the three lineages, syngnathid, cypriniform, and perciform, from our study (i.e., putative gene losses or highly derived genes). Using the 28 species sampled in the GENESPACE analysis, we determined a list of genes that were not detected in each clade, present in the closest sampled outgroup, and present in the two other scRNAseq representatives. For example, genes present in mandarin dragonet (outgroup), stickleback and zebrafish, but undetected in all 5 sampled syngnathids. These putative losses include 170 genes in syngnathids, 1 gene in cypriniforms, and 3 genes in perciforms (Fig. S4). The list of genes not identified in syngnathids includes published losses (*odam, fgf3, fgf4, fgf9*, *eve1*, and *tbx4)*. Although the additional losses may be relevant for syngnathid craniofacial evolution – for example *elk4* and *cacna1ba*, genes in the MAPK signaling KEGG pathway - many candidates have broad expression patterns and have no known connection to craniofacial development.

To identify highly derived and novel genes we next used the GENESPACE analysis to distinguish orthologs present in pipefish, zebrafish, and stickleback that were found in all members of their lineage but with no detected orthologs or genes with identical names in the other sampled species. We found 26 genes with orthologs present in all syngnathids but not detected in other species, 174 zebrafish genes uniquely detected in all cypriniforms, and 0 stickleback genes with specific detection in sampled perciforms. Since the number of unique genes depends on the evolutionary divergence of species included in the GENESPACE run, we focused interpretation on gene expression patterns rather than gene quantity comparisons. The genes uniquely detected in syngnathids had weak expression in the atlas, not specific to craniofacial cell types (Fig. S5). Similarly, no genes with high pharyngeal arch expression were observed in the cypriniform list (Fig. S6). Overall, we found no evidence in these data for the evolution of novel genes specific to syngnathid craniofacial gene networks.

Finally, we investigated whether the evolution of regulatory elements could impact craniofacial development in syngnathids by using a whole genome alignment (WGA; 12 species) and ATAC-seq data (zebrafish, stickleback, and Gulf pipefish) paired with the scRNAseq atlas. Since single cell libraries are prone to technical noise (dropout), we focused on changes in gene expression presence or absence to evaluate the potential of a conserved element (CE) to relate to gene expression rather than differential abundance. We found 24 genes with CE unique to syngnathids included in the WGA and with unique Gulf pipefish expression for any scRNAseq cluster. These genes included several collagens (*col17a1b*, *col8a2*, and *col9a2*) and the candidate *ece1,* which interacts with the endothelian pathway and is predicted to act upstream of viscerocranium morphogenesis (Fig. 4A; 25). Notably, the syngnathid specific CE in *ece1* overlaps with an ATAC-seq peak from Gulf pipefish and is within the 5’ UTR of the gene (Fig. 4B). Although *ece1* unique expression was also observed in a neural cluster (#5), a greater percentage of Gulf pipefish CNC PA cells (25%) express this gene than do the PA cells of the other two species (Gac 14%, Dre 4%).

**Fig. 4.**
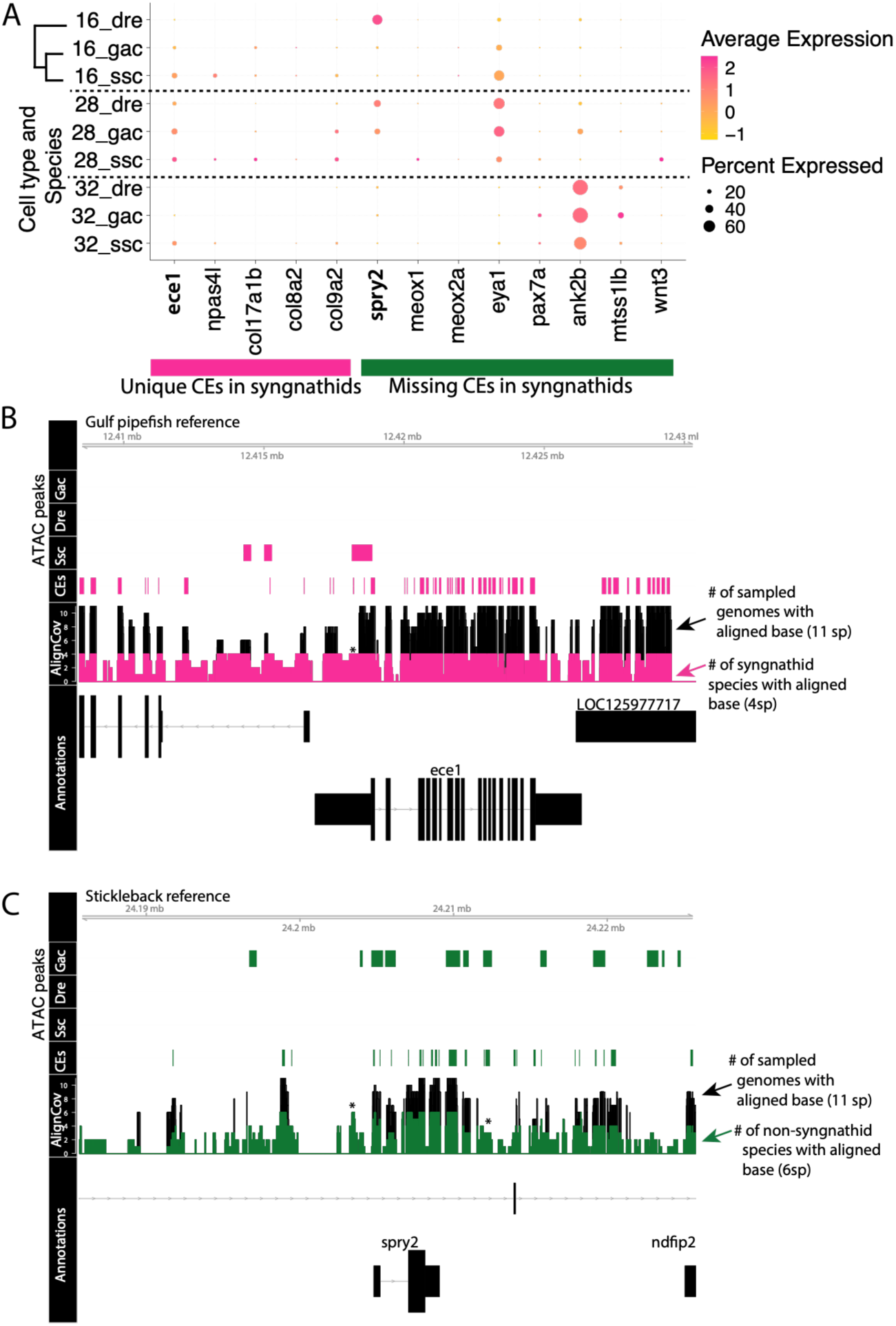
Genes with syngnathid-specific changes in conserved elements also have expression differences. Expression of genes with unique conserved elements in syngnathids (pink) and missing conserved elements (CEs) in syngnathids (green) are displayed in a dot plot (panel A). Expression from cluster 16 (CNC PA cells), cluster 28 (pharyngeal pouch cells), and cluster 32 (neural CNC-derived cells), shown in the plot. Using the Gulf pipefish genome as a reference, conserved elements from pipefish (CEs), ATAC-seq, and whole genome alignment (AlignCov) data are plotted for *ece1* (exon structure depicted at bottom) (panel B). Whole genome alignment shows the alignment coverage for each base on the Gulf pipefish genome, with syngnathid specific coverage highlighted in pink. Using the threespine stickleback genome as reference, conserved elements from stickleback (CEs), ATAC-seq, and whole genome coverage (AlignCov) data are shown for *spry2* (panel C). Coverage data captures the number of genomes that aligned to each base of the stickleback genome, with non-syngnathid genomes highlighted in green.

In addition to genes with syngnathid-specific CEs identified in the WGA, we also identified 180 genes with CEs present in all species except for sampled syngnathids.

This list included developmental genes such as *meox1, meox2a, eya1*, and *pax7a*. Since none of these candidates had expression unique to a “short snout” (stickleback and zebrafish) cell cluster, we also identified CEs that are present in all the analyzed percomorph species except syngnathids and that had expression unique to any zebrafish or stickleback cluster. This analysis revealed 4 candidates, *ank2b*, *mtss1lb*, *spry2*, and *wnt3* (Fig. 4A). In *spry2* there are several CEs not conserved in syngnathids, including an element that overlaps with an ATAC-seq peak from stickleback (Fig. 4C).

Intriguingly, *spry2* is a negative regulator of the Fgf signaling, such as in repressing tooth development by repressing sensitivity to signaling by *fgf3* (26) and also has a role in facial outgrowth that could be independent of FGF signaling (27).

### Novel Fgf expression in Gulf pipefish pharyngeal arch cells

With evidence for changes in gene and CE content of signaling pathways, we compared expression and examined CEs for ligands, receptors, and interactors from key gene families for head development (Fgf, Bmp, Notch, Endothelin, and Wnt) across zebrafish, stickleback, and Gulf pipefish cells (Fig. S7-12). While the presence or absence of signaling gene expression in CNC derived PA cells was largely conserved among all three species, we found select cases of unique expression loss and gain in Gulf pipefish. Genes previously identified as missing in syngnathid genomes (*fgf3* and *fgf4)* had PA cell expression in zebrafish and stickleback CNC derived cells and pouch cells (Fig. 5A). Although *fgf3* and *4* are still present in the bluespotted cornetfish, which is an outgroup to the syngnathid clade, cornetfish *fgf4* did not align with *fgf4* genes of other species in the WGA and two small *fgf3* CEs from exons 2 and 3 were not found (Fig. S13A). Potentially, this indicates sequence divergence or degradation predating the tandem loss of these two genes in syngnathids.

**Fig. 5.**
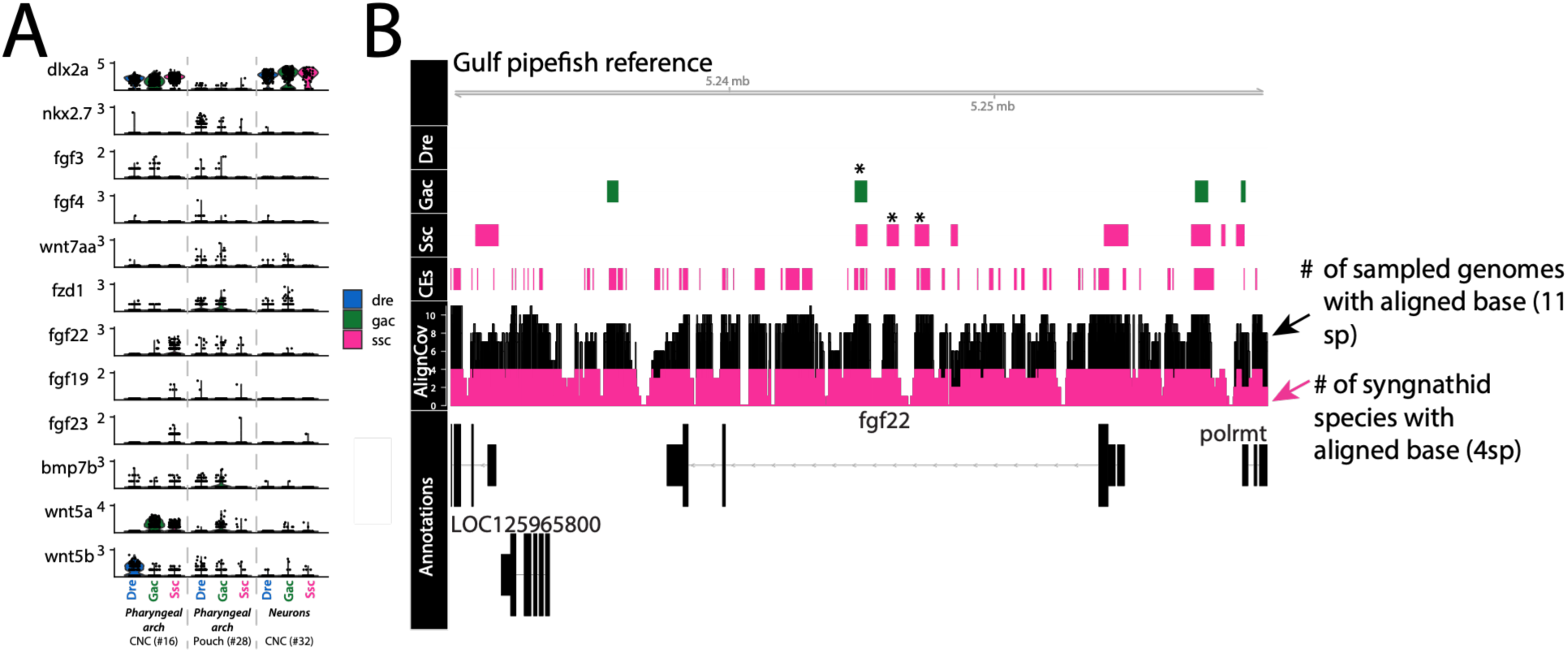
Novel signaling pathway expression changes in Gulf pipefish CNC pharyngeal arch cells and changes in potential regulatory elements in Percomorphs. Expression pattern of signaling genes is displayed in a Violin plot in which each dot is a cell and species are plotted individually (panel A). The genes *dlx2a* and *nkx2.7* are canonical markers of CNC derived cells (clusters 16 and 32) and pharyngeal arch epidermal cells (cluster 28), respectively. Selected signaling genes have expression differences between species. The genomic landscape of *fgf22*, a gene with percomorph specific CNC expression, is represented in panel B by the pattern of Gulf pipefish conserved elements (CEs), ATAC-seq peaks, and whole genome alignment coverage with syngnathid species coverage in pink.

From our GENESPACE analysis, two Wnt signaling pathway genes, *fzd1* and *wnt7aa*, were not detected within the sampled syngnathid lineages. These genes were expressed in PA endodermal pouch cells in stickleback and zebrafish, and *fzd1* was also expressed in CNC-derived PA cells. Although neither gene was recovered in the orthology search, whole genome alignment coverage data shows partial alignment of syngnathids genomes of both genes, suggesting that they are highly derived or pseudogenes (Fig. S13B).

We observed intriguing patterns in four genes of those present in all three species: *fgf22, fgf23, fgf19*, and *bmp7b.* All three *fgf* genes were absent in zebrafish CNC derived PA cells in the scRNAseq analysis, but present in pipefish. While stickleback also lack CNC PA expression of *fgf23* and *fgf19*, *fgf22* was minimally expressed in PA cells. Specific PA expression of *fgf22* in stickleback and pipefish was confirmed using *in situ* hybridization, and an additional novel domain of expression was observed in stickleback fin buds (Fig. S14). For *fgf22,* there was a high degree of sequence conservation between species (excluding zebrafish; Fig. 5B). These conserved elements aligned with ATAC-seq peaks from stickleback and pipefish, suggesting a functional relevance. Despite presence in all species except for *D. excisus*, syngnathid *fgf19* and *fgf23* sequences did not align with non-syngnathid species (Fig. S13). These misalignments are suggestive of genomic rearrangements such as rearrangements of syntenic genes at the fgf19 locus previously reported in several syngnathids (28). Interestingly, *fgf19* in the ancestral genomic condition is adjacent to the syngnathid-missing genes *fgf3/4* and *fgf23* ancestrally neighbors *tigarb,* which is lost in most syngnathids. Genic rearrangements such as segmental deletion could have led to changes in expression of syngnathid *fgf19* and *fgf23* by mechanisms such co-option or loss of regulatory sequences.

Outside the Fgf family, *bmp7b* was expressed in CNC derived PA cells of all species, but uniquely absent in the PA pouch of Gulf pipefish. In the case of *bmp7b,* Gulf pipefish lacked expression in the PA pouch, and there was a CE missing in the first intron of *bmp7b* in most syngnathids except *Doryrhamphus excisus.* This CE, however, was not strengthened as a candidate functional element by overlap with ATAC-seq peaks (Fig. S15).

In addition to syngnathid specific findings, we found deviations in the two percomorphs vs. zebrafish in their expression of signaling genes. Primarily, these changes were seen in paralogs derived from the teleost whole genome duplication event. We found differential gene expression of *wnt5* paralogs, *wnt5a* and *wnt5b* (Fig. 5A). Pipefish and stickleback expressed both paralogs in CNC-derived PA cells, but zebrafish expressed only *wnt5b* in these cells. We also found evidence for differential gene retention of paralogs created by the teleost whole genome duplication. Zebrafish retained *fgf18b*, *jag2a*, *fzd8b, dkk1b, bmp2a, bmpr1ab, bmpr2b, sfrp1b, nkd2b,* and *fgfbp2a* but these genes have not been identified in stickleback or pipefish genomes (Fig. S7-12). Although these findings may not directly relate to the unique craniofacial morphology of syngnathids, the genetic and genomic changes establish potentially significant lineage-specific differences between percomorphs and zebrafish CNC GRNs that should be considered when extrapolating inferences from experimental biology based in the zebrafish model.

### Increased number of CNC PA cells warrants future investigations

Apart from the somitic mesoderm, which was congruent across the compared lineages, the three species varied in their cellular contribution to clusters (Fig. 6). We suggest that this variance may originate from tissue dissociation or biological differences rather than indicating major developmental stage mismatch. We found that species largely had congruent cellular differentiation states suggesting that these datasets are comparable (Fig. S16; 27). Additionally, cell number disparities spanned cell lineages that are not well explained by cells being in different developmental stages (such as increased zebrafish cells from the epidermis/periderm).

**Fig. 6.**
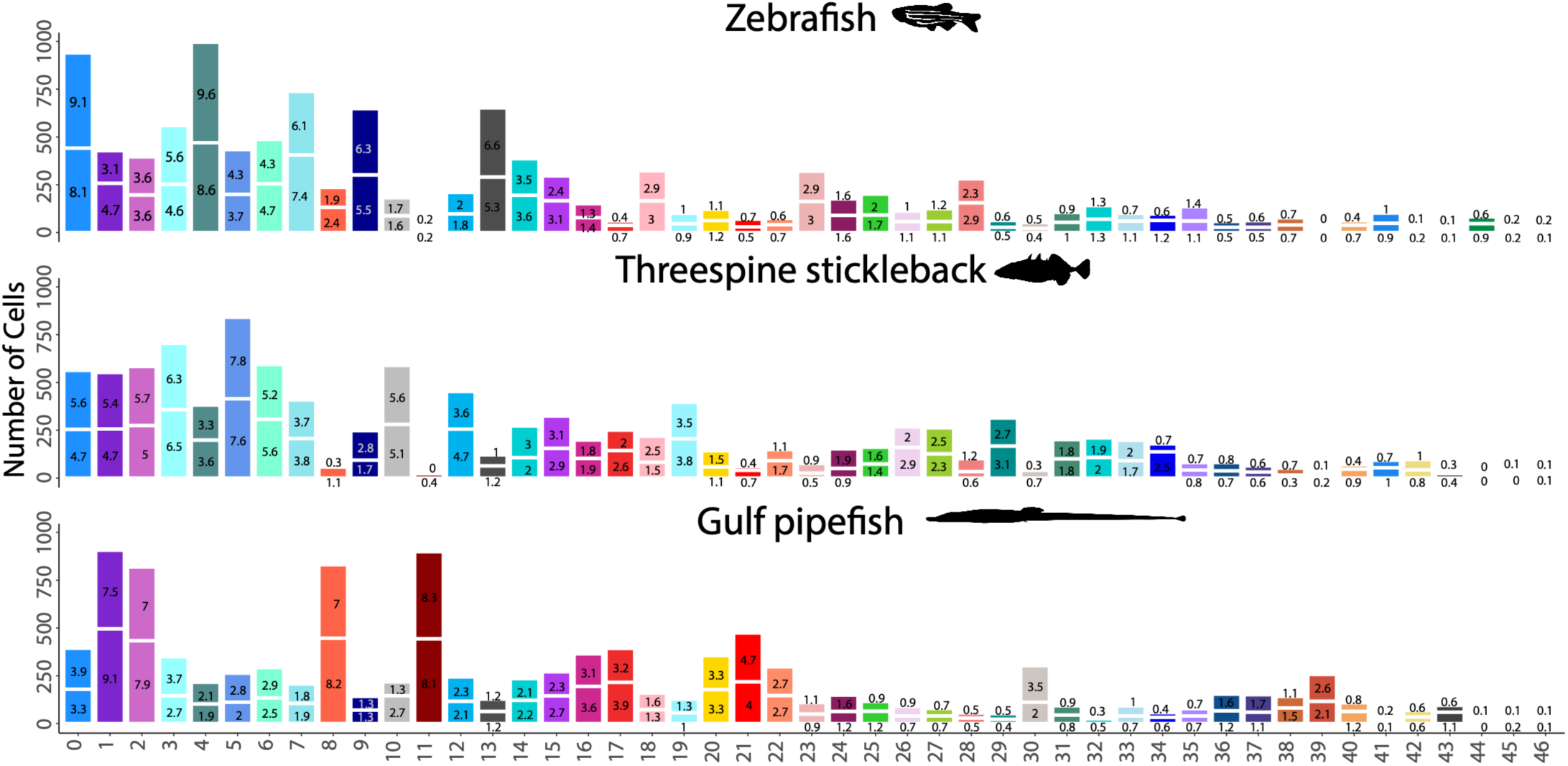
The number of cells per cluster varied by species. Species data are separated into three panels with the two replicate samples distinguished by a white line. The percentage that each cluster contributes to the sample’s cell total is labeled on the bars.

It is possible that differences in biology (e.g. cell proliferation/death) and methodology (e.g., ease and success of dissociating different cell types) can explain these cell number differences. Most strikingly, erythrocyte cells contribute to roughly 20% of the pipefish cells but around 2% of stickleback and zebrafish samples. This extreme difference contributed to poor integration of the RBC clusters. Although pipefish embryos may have increased numbers of RBC cells, it is also possible that these cells may have dissociated more easily or originate from the well vascularized pseudoplacenta of the father’s brood pouch.

A noticeable difference in Gulf pipefish cells was confidently inferred as intrinsic to the embryos is in CNC PA cells. These cells accounted for 3.35% of all Gulf pipefish cells, nearly double their contributions in stickleback (1.85%) and zebrafish (1.35%). To probe whether this change in PA cells could relate to a higher overall number of CNC cells of various fates, or a greater proportion of CNC cells differentiating specifically into skeletogenic cells, we compared PA and neural cells derived from the CNC. We found that stickleback and pipefish had roughly the same number of total CNC cells (stickleback = 403; pipefish = 409), but that pipefish had a higher relative number of PA cells (stickleback = 199, pipefish = 365) than neural cells (stickleback = 204, pipefish = 44). Zebrafish, like stickleback, had relatively equal numbers of PA and CNC-derived neural cells (141 neural cells; 151 PA cells). This finding provides preliminary evidence for differential cell abundance of PA cells in Gulf pipefish and motivates detailed studies comparing PA cell numbers across species.

## Discussion

In this study, we created the first comparative, developmental scRNAseq atlas to determine the scale of GRN re-wiring in syngnathid craniofacial development. We showed that Gulf pipefish CNC cells have not experienced global GRN re-wiring despite having highly derived head and jaws and numerous gene losses. Although we found gene turnover in syngnathids with 26 potential novel genes and 170 suspected gene losses, most of these candidates were not expressed in CNC cells or were likely auxiliary to the core network based on their broad expression. However, local GRN changes, including strong *fgf22* expression and co-option of *fgf19* and *fgf23* in CNC cells, and changes to potential regulatory elements in *ece1* and *spry2,* might explain the resilience of syngnathid development to the loss of *fgf3*, and could relate to elongated snouts. *fgf22* and *fgf3* are both members of the mitogenic, “Fgf7 subfamily” of the Fgf ligand genes, and both ligands share affinity to FGFR1b and FGFR2b receptors (reviewed in 28, 29). We propose that expression of *fgf22* in syngnathid CNC cells provided redundancy with *fgf3*, permitting *fgf3* loss. A co-option of the role of *fgf3* by *fgf22* could have resulted in increased CNC proliferation if *fgf22* is upregulated in syngnathids as our RNAseq and *in situ* data suggest. This hypothesis assumes zebrafish *fgf22* expression pattern represents the ancestral state, it is possible that zebrafish have lost *fgf22* PA expression . Overall, our analyses support the hypothesis that pathway redundancy enabled gene loss and phenotype evolution occurred without widespread network changes in syngnathids. This case study of GRN evolution showcased the conservation of core gene regulatory networks in an organism with conspicuously derived heads, while instead a few key changes could have led to this phenotypic evolution.

### No evidence for global re-wiring in pipefish CNC

Through examinations of developmental pathways, gene losses, potential novel genes, and gene networks, we found a small number of candidate genes for the evolution of syngnathid derived faces. In addition to Fgf pathway changes, we identified degenerated genes in the Wnt (*fzd1* and *wnt7aa*) and an expression domain loss in the Bmp (*bmp7b*) pathway. These genes are all expressed in the pharyngeal arch. Although there is no evidence to suggest their individual losses would impact craniofacial morphology (32–35), the possible consequences of their combined losses is currently unexplored through experimental manipulation in models such as zebrafish. Extending from known signaling pathways to *de novo* candidates, we found evidence for gene turnover in syngnathids but did not identify suggestive candidates for craniofacial evolution. Our regulatory element analysis found new conserved elements in *ece1,* a metalloprotease that processes the endothelin-1 receptor protein for the endothelin pathway, and *npas4l,* a transcription factor downstream of Bmp signaling. We also found increased expression of these genes in pipefish CNC cells (36, 37). Loss of *ece1* mRNA in mice and zebrafish leads to drastic craniofacial defects, with changes to dorsal and ventral elements in zebrafish, suggesting changes to this gene could impact pipefish craniofacial development (25, 38). Consistent with examinations from other species (birds, cichlids), these analyses suggest local rather than global re-wiring drives syngnathid craniofacial evolution (4, 39).

### Differential fate of teleost whole genome duplication paralogs leaves a strong signal in developmental expression data

In comparison to the minimal GRN changes between syngnathids and “short” snouted fishes, we observed a strong phylogenetic signal in the expression of some genes, particularly paralogs created by the TGD. We identified 11 signaling pathway gene losses and 2 expression domain differences in percomorphs vs. zebrafish TGD paralogs. Expression discrepancy was observed in highly conserved craniofacial development genes *ednraa* and *ednrab*. These genes are partially redundant in zebrafish, though fish with knockdowns of *ednrab* have shortened lower jaws and joint fusions (40). In contrast, our analysis found that percomorph members did not express *ednrab* in pharyngeal arch cells. This incongruity suggests differential subfunction partitioning between paralogs in the two fish lineages (41). Beyond the biological finding, this observation highlights that the loss of *ednrab* expression in Gulf pipefish PA cells could have been mis-interpreted as developmentally relevant if threespine stickleback had not been included in the analysis. This finding argues for comprehensive inclusion of species in future studies focused on making inferences about GRN evolution.

Although our analysis did not find evidence for global GRN re-wiring in syngnathids, the observed phylogenetic signal implicates developmental systems drift (42). Under this principle, developmental genetic mechanisms can change over evolutionary time, though phenotypic outcomes for homologous structures do not. Therefore, teasing out genetic changes unrelated to morphological evolution is critical for removing false positives from evo-devo studies. Further, it indicates that cross-species scRNAseq is challenging to interpret without adequate phylogenetic sampling.

### Surprising pipefish expression of *fgf23* calls into question its developmental role

Fgf signaling constitutes highly conserved pathways active in craniofacial, somite, limb, and neural development in vertebrates (30). Our dataset revealed the Gulf pipefish endocrine Fgf ligands (primarily *fgf23)* surprisingly deviate from this this pattern of conservation. Although *fgf23* in model systems is involved with the regulation of calcium and phosphate metabolism by kidneys and bone (43–45), we observed *fgf23* expression in pipefish PA CNC, mesenchymal, somite, and retinal cells. The sequence of *fgf23* is conserved in syngnathids, suggesting maintenance of binding affinities, but loss of neighboring gene *tigarb* around could have altered the regulation of *fgf23* and led to these novel expression domains. If this hypothesis is true, we predict the basal syngnathid *Doryrhamphus excisus* could lack *fgf23* PA expression because the species has retained *tigarb*.

Novel *fgf23* expression domains may represent a shift from endocrine to paracrine signaling. This gene is suspected to engage in paracrine or autocrine signaling in mouse osteoblasts due to expression of *fgf23* and its receptor complex (*alpha-klotho* and *fgfr1;* 46). However, our dataset found minimal klotho expression (<15% of cells across all clusters), suggesting pipefish arch *fgf23* signal transduction does not occur via this canonical mechanism. An alternative signaling mechanism for *fgf23* has been identified in rat cardiac myocyte cells where phospholipase Cy acts as a co-factor and the calcineurin/NFAT pathway is activated. (47). Since phospholipase C (plc) genes are expressed in arch and mesenchymal cells, with *plcd3a* and *plcd4b* unique to pipefish arches, it is possible that *fgf23* is signaling through this alternative pathway in developing pipefish. Although the role of *fgf23* in pipefish development is unknown, it is a noteworthy case of possible signaling gene co-option in GRNs of the PA, somites, and mesenchyme.

### Evolution of Fgf pathway could have created resiliency to gene loss and impacted craniofacial development

Our study proposed losses and gains in Fgf pathway redundancy in cypriniform and percomorph fishes are relevant for syngnathid evolution. Considering syngnathids’ loss of *fgf3*, we examined the other members of its Fgf subfamily (*fgf3*, *fgf7*, *fgf10*, and *fgf* 22; 28) and detected PA expression of fgf*10a, -10b*, and -*22* in stickleback, fgf*10a, - 10b*, and *-22* in pipefish, and *fgf10a* and -*10b* in zebrafish. Based on inferred ancestral expression of *fgf3* (CNC among other domains) and *fgf22* (neural), it is parsimonious to hypothesize that *fgf22* gained expression in percomorph PA cells and developing fins. In this case of local GRN re-wiring, *fgf3* may have become redundant in percomorph PA CNC cells with *fgf22* mitigating consequences of losing *fgf3* in syngnathids.

While percomorph *fgf22* lends an apparent gain in redundancy in the Fgf pathway, we suggest that percomorph *fgfr1* paralogs have decreased in redundancy. The gene *fgfr1* has two paralogs in teleost fishes, which each have two splice variants (1b and 1c) that vary in Fgf ligand affinity. We observed that percomorphs are missing the exon for the 1b splice variant in *fgfr1b*, and syngnathid fishes are missing the *fgfr1b* gene completely (28; Fig. S17-18). Although *fgfr1a* and *fgfr1b* are redundant in zebrafish, the loss of *fgfr1a* expression in medaka is lethal (48, 49). The loss of the *1b* splice variant in *fgfr1b* likely leads to reduced binding of *fgf3*, *7*, and *8* by *fgfr1b*, possibly explaining the detrimental effects of losing *fgfr1a* expression in medaka (50). Combined, these evolutionary (syngnathids) and experimental (zebrafish and medaka) knockouts suggest varying redundancy in the Fgf signaling pathway across teleosts.

Redundancy in GRNs increases robustness against perturbations, illustrated by the maintenance of PA 3-7 in syngnathids despite *fgf3* loss (51). An interplay between developmental robustness and evolvability (52) exists, however, and exploring this interplay with *fgf22* presents relevant hypotheses for syngnathid craniofacial evolution. In the zebrafish, *fgf22* knockdowns experience reduced cell proliferation in the midbrain (53). We propose *fgf22* may have a similar role in proliferation in stickleback and pipefish pharyngeal arches and fins. Indeed, *fgf22* is expressed in the proliferative mesenchyme proximal to the apical ectodermal ridge in stickleback and pipefish pectoral fins. This expression mirrors the fin domain of *fgf10a*, suggesting co-option, robustness, and a similar role regulating limb outgrowth (54, 55).

Although *fgf22* expression was weakly detected in stickleback pharyngeal arches, in Gulf pipefish we observed strong expression in the gill arches and around the periphery of the first and second arches. scRNAseq revealed that Gulf pipefish abundance of fgf22-expressing CNC PA cells and average *fgf22* expression were higher compared to either *fgf22* or *fgf3* in stickleback. We found that Gulf pipefish had a greater number of CNC derived PA cells than stickleback and zebrafish, suggesting the possibility that increased fgf22 expression could result in greater cell proliferation in the gill arches. Although it is possible that differences in detected cell numbers result from technical differences in scRNAseq dissociations, the suggestive findings warrant subsequent study. Furthermore, we found a loss of conserved noncoding elements linked to *spry2* and altered expression of this key negative regulator of the Fgf pathway, presenting a possibility that reduced expression of *spry2* enabled increased Fgf responsiveness to signaling ligands (i.e., *fgf22* and others; 56). Experimentally teasing out the roles of *fgf22* and *spry2* on CNC proliferation in tractable percomorph models such as medaka or stickleback could provide important insight into the unique morphology, particularly in the highly derived second arch, of syngnathids (57, 58).

## Materials and Methods

### Production of scRNAseq sequencing libraries

We created single cell RNA sequencing atlases from replicates consisting of two pools of embryos from each species, threespine stickleback and Gulf pipefish. Samples were sequenced at the onset of pigment development in the retina (70hpf for stickleback and around 5-6 dpf for pipefish). For stickleback 70hpf sample one, we used the scRNAseq data from (59). For stickleback 70hpf sample two, twenty-seven 70hpf stickleback embryos were dissociated in 0.25% trypsin as per (59) and filtered with a 40µM cell strainer (MTC Bio C4040). Pipefish samples 1 and 2 came from a 5dpf and 6dpf clutch, respectively. Fathers were euthanized using MS-222, embryos were removed from the pouch then dissociated. 20 pipefish embryos per sample were dissociated using 460 µL TrypLE Express (Gibco #12605-010) and 40 µL collagenase I (sigma C0120-100MG) for 8.5 minutes, filtered through 40µm cell strainer, quantified with a Bio-Rad cell counter, diluted to a concentration 800 cells/uL. All samples were prepared with the 10X Genomics Single Cell 3’ v3.1 Dual Index Genome Expression (GEX) – mRNA-Seq kit at the Genomics and Cell Characterization Core Facility (GC3F) at the University of Oregon. Library sequencing was performed on an Illumina NovaSeq 6000 platform using paired-end 150 base-pair reads. Sequencing yielded 864 and 901 million reads in stickleback and 766 million and 1.2 billion reads in pipefish.

### Production of ATACseq sequencing libraries

ATAC-seq libraries were produced from threespine stickleback and Gulf pipefish at the onset of retinal pigmentation. For Gulf pipefish, dissociated cells from the scRNAseq libraries were used as the input to the ATACseq library prep kit. For threespine stickleback, we dissociated cells from 15-20 70 hpf embryos using TrypLE Express. 340,000 cells were used as the input to the Active Motif kit (part # 105320) to prepare ATACseq libraries. Sequencing was performed on an Illumina NovaSeq 6000 platform using paired-end 150 base-pair reads. Sequencing yielded 24 and 59 million reads for stickleback samples and 43 and 328 million reads for pipefish samples.

### scRNAseq library initial processing

Two zebrafish 24hpf (60) scRNAseq datasets were downloaded using SRA toolkit (v3.0.7). The scRNAseq reads for all species were processed using CellRanger (v7.2). For zebrafish, we aligned reads to the Lawson genome assembly (GRCz11_4.3.2/danRer11) and annotation (v4.3.2; 62). For stickleback, we aligned reads to the GCF_016920845.1 assembly with its respective annotations extended with scISOseq reads (reads from 60). We aligned Gulf pipefish scRNAseq reads to the GCF_024217435.1 genome and annotation file from (62). Using the CellRanger outputs, we identified doublets using scrublet.

### Ortholog identification

We identified orthologs between zebrafish, pipefish, and stickleback using GENESPACE (63). We used a variety of species to identify orthologs - from solely the three focal species to the inclusion of 27 additional, high-quality genome assemblies - to find the highest number of 1:1:1 orthologs (Supplementary file). We also assembled a test set of genes from stickleback and zebrafish to test for accuracy in ortholog identification (Supplementary file). After finding the highest number of 1:1:1 orthologs and accuracy of ortholog identification, we chose to use our GENESPACE analysis with 3 species and 25 additional species. From the GENESPACE pangenes output (1 per species per run), we isolated rows from the dataset where the orthogroup column contained stickleback, zebrafish, or pipefish, merged based on common orthogroups, and wrote out csv files using custom scripts (Code is available on Github, https://github.com/hopehealey/cross_species_scRNAseq_analysis). Then, merged the 3 cleaned pangenes files per run, removed any duplicates, and output the final pangenes csv. We used the final pangenes csv file to rename genes from the Cell Ranger outputs to enable scRNA-seq dataset integration. Using custom scripts, gene names from the filtered_feature_bc_matrix output files were modified according to orthology relationships (Table S3). Because zebrafish is a model system with available resources and databases, orthologs were by default changed to zebrafish names (and thus, zebrafish filtered_feature_bc_matrix output files were not modified). For stickleback, gene names were changed to zebrafish names if the gene had a 1:1 ortholog with zebrafish. For pipefish, gene names were changed to zebrafish names if the gene had a 1:1 ortholog with zebrafish. Additionally, if a pipefish gene did not have a 1:1 ortholog with zebrafish but it did have one for stickleback, gene names were changed to the stickleback name. For 1:many and 1:0:0 orthology cases, genes were not renamed.

### scRNAseq atlas construction

Using Seurat (v5.0.3), we pre-processed all the datasets separately. We removed ambient reads with SoupX (v1.6.2) and doublets with scrublet. Based on the distribution of samples’ read counts per cell, we filtered the datasets to remove cells with high numbers of reads. Cells with fewer than 100,000 counts in stickleback and pipefish, and fewer than 40,000 counts in zebrafish, were retained. We also removed cells with low (< 500 genes) and high numbers of detected genes (> 10,000 detected genes in stickleback and pipefish and > 6,000 genes in zebrafish), and high percentage of mitochondrial genes (> 5% of mitochondrial reads). Finally, samples were randomly down sampled to the size of the smallest scRNAseq atlas (pipefish 5dpf; 5,469 cells).

The six equally sized datasets were integrated using Seurat Canonical Correlation Analysis (CCA) method. The integrated dataset was clustered with 40 PCs at a 1.9 clustering resolution. Integration was tested with a varying number of PCs (from 11 to 50), and 40 was chosen because it correctly resolved the two expected neural crest clusters with the fewest PCs. The success of integration was evaluated using LISI (v1.0). Marker genes were identified using FindAllMarkers and FindConservedMarkers from Seurat. To annotate cluster identity, we examined marker gene expression from ZFIN and Daniocell, then proposed identity based on several markers for each cluster (18–21). We further examined marker gene expression in different species using DotPlots. We visualized gene expression patterns, cell abundances, and cell distribution in multidimensional space using Seurat, ggplot2 (v3.5.1), and scCustomize (v2.1.2).

### scRNAseq trajectory analysis

To identify whether species’ samples were at similar developmental stages, we predicted differentiation status by species using CytoTRACE (v0.3.3(29). For each cluster, we randomly sampled 125 cells from each species, extracted the counts layer, and used iCytotrace to identify differentiation status.

### scRNAseq differentially expressed gene analysis

To identify genes that varied in expression level across species, we performed differential expression analysis on each cluster individually. Since the number of cells per cluster varied by species, we randomly downsampled each species’ cells to 100 cells. For each cluster, we ran FindMarkers in three pairwise comparisons (stickleback vs. pipefish; pipefish vs. zebrafish; zebrafish vs. stickleback). Next, we examined these data in different subsets: genes expressed in pipefish but not in the other fish, genes expressed in stickleback but not the other fish, genes expressed in zebrafish but not the other fish, genes expressed in both pipefish and stickleback but not zebrafish, genes expressed in both zebrafish and pipefish but not stickleback, and genes expressed in both stickleback and zebrafish but not pipefish. Genes were then filtered to remove genes with adjusted p-values above .05 and logFCs below 1. Finally, genes were removed from the analysis if they were not 1:1:1 orthologs (i.e. no lost or potentially novel genes were retained).

### scRNAseq gene network analysis

To discover derived GRNs in Gulf pipefish we built genetic networks using WGCNA (v1.72-5; (24). First, we built gene networks from the entire atlas and species-specific derivations of the atlas. For each approach, we normalized the dataset, ran FindVariableFeatures to identify 2,000 variable features, and scaled the data. Next, we created gene expression matrices using each dataset’s variable genes, picked an appropriate soft threshold using pickSoftThreshold (pipefish=3, stickleback=5, zebrafish=2, entire dataset=2), transformed the dataset with the soft threshold, and created a dissimilarity matrix. To build gene networks, we then clustered genes using hclust, created modules with cutreeDynamic (minimum cluster size of 15, guide hang of .05, and hang of .03), and calculated module eigengenes and merged modules with a module eigenegene correlation of 70% or higher. We determined the correlation of each module with cell clusters by calculating the mean correlation of module eigengenes of each cell in the cluster.

We evaluated whether each cell cluster impacted network structure by completing a connectivity analysis. We calculated the baseline connectivity for each module, deriving the mean connectivity inside and outside the module. Then, each cell cluster was progressively removed and the connectivity was re-measured. To determine whether the change in connectivity was significant, we completed 1,000 permutations where cells (equal to the size of the cluster) were randomly removed from the analysis-wide dataset and then measured the connectivity again. We determined p-values by comparing the number of cases where the change in connectivity was higher in the permutations than the removal of the cell cluster. Lastly, p-values were corrected with FDR.

### scRNAseq cross-species correlation analysis

To assess whether Gulf pipefish have divergence in overall expression profiles, we completed a correlation analysis comparing species in a pairwise manner. Using code derived from (23), specifically compare_nhoods.R, we built neighbors of cells for each species using miloR (v1.8.1; k = 40, d=40, prop=.3, using the CCA integrated dimension). We then calculated neighborhood correlations between every neighborhood in the two species using 8,000 highly variable genes which were filtered to include only 1:1:1 genes. To compare clusters across species, correlation scores were filtered to ensure that the index cell of compared neighborhoods belonged to the same cluster.

We completed this cluster level comparison with binned (combining clusters with similar annotations) and un-binned (analyzing each cluster individually) cell clusters. For the binned cell cluster approach, we created 11 categories: pharyngeal arch CNC cells (cluster 16); mesenchymal, somites, and notochord cells (1, 2,15, and 35); unknown cells (10); neural cells (0,3,4,5,6,7,9,12,13,14,19,29,31,32,33,34,36,37,41,43, and 46); red blood cells (8,11,17,21,22, and 39); endothelial cells (24); epithelial cells (18, 23, 26, and 28); muscle cells (25, 27, 44, and 45); immune cells (30); pigment cells (38 and 40); and endodermal cells (20 and 42).

### ATACseq read processing and analysis

We downloaded two samples of 24hpf zebrafish ATAC-seq data from (64) using SRA toolkit (3.0.7). Reads from stickleback, pipefish, and zebrafish were aligned to their respective genome assemblies using Bowtie2 (v2.3.4). Mitochondrial reads were then removed from the threespine stickleback and zebrafish genome assemblies using grep. The Gulf pipefish reference genome does not include mitochondrial sequences. Reads were filtered to remove those with low quality scores (q10) and sorted using samtools (v1.19). Next, we marked and removed duplicate reads with picard (v2.6.0). We then sorted reads by name using samtools (v1.19) to prepare them for peak identification using genrich (v0.6.1) with both sample replicates.

### Whole genome alignment and identification of conserved elements

To determine whether genes with detected expression changes in Gulf pipefish may have alterations in regulatory elements, we paired our ATAC-seq analysis with a conserved element approach. First, we selected 12 species: 5 syngnathids, comprising Gulf pipefish (*Syngnathus scovelli*), dwarf seahorse (*Hippocampus zosterae*), leafy seadragon (*Phycodorus eques*), scribbled pipefish (*Corythoichthys intestinalis*), and bluestripe pipefish (*Doryrhamphus excisus*); 1 long-snouted outgroup, bluespotted cornetfish (*Fistularia commersonii*); 1 outgroup with a “short” snout, mandarin dragonet (*Synchiropus splendidus*); 5 additional species, comprising zebrafish (*Danio rerio*), northern pike (*Esox lucius*), Japanese medaka (*Oryzias latipes*), threespine stickleback (*Gasterosteus aculeatus*), and southern bluefin tuna (*Thunnus maccoyii*).

We identified repeat elements with RepeatModeler (v 2.0.5) and masked them with RepeatMasker (v 4.1.5). For RepeatMasker, we used concatenated repeat files generated from all species RepeatModeler results as the repeat library input. Masked genome assemblies were aligned using Cactus (v2.8.1; 67). We used halAlignmentDepth (v2.2) to identify number of genomes that aligned respective to a reference (--noAncestors) using the –targetGenomes parameter to select subsets.

Then, for each chromosome of each species, we generated maf files using cactus-hal2maf and identified conserved elements using phastcons (v1.5; target-coverage 0.125, expected-length 12, most-conserved). Conserved element bed files were concatenated to include all chromosomes for each species separately. Chain files were then generated using cactus-hal2chains to create pairwise species maps. Finally, conserved elements and ATAC-seq peaks were lifted over using these chain files to respective genome assemblies using UCSC liftover (minMatch=.75). Elements and ATAC-seq peaks were visualized using IGV (v2.16.2) and Gviz (v1.56.0).

### *In situ* hybridization

For the gene *fgf22*, we completed *in situ* hybridizations in Gulf pipefish, stickleback, and zebrafish. Primer sequences were designed using NCBI Primer Blast and synthesized embryonic cDNA libraries. Probes were prepared using TOPO cloning. To procure Gulf pipefish embryos, we allowed the fish to mate in our aquatic rearing facility then harvested embryos once they reached the appropriate stages. Gulf pipefish were reared in 25° C water with 25-28 ppt salinity. To gather stickleback embryos, we completed *in vitro* fertilization with fish from a line originally derived from a wild population collected from Cushman Slough (Oregon, USA). Embryos were reared at 20° C using standard stickleback rearing procedures (66) and harvested at necessary stages. Zebrafish were obtained from the Aquatic Animal Care Services (AqACS) at the University of Oregon at 24 hpf. Embryos and larvae were fixed in 4% PFA, dehydrated through a series of PBT/MeOH washes, and stored in MeOH at -20° C. 5-12 embryos were used for each experiment. We completed *in situ* hybridizations in keeping with (67), leaving the embryos in stain until background was observed. After imaging, we used the levels tool in Adobe Photoshop (v23.4.2) to white balance the photographs.

## Data accessibility

All raw sequencing files generated in this publication have been uploaded to NCBI (PRJNA1420934). Updates to the Gulf pipefish and stickleback annotation files will be published on Dryad. Custom scripts are hosted on Github (https://github.com/hopehealey/cross_species_scRNAseq_analysis).

## Acknowledgements

We thank Mark Currey for culturing the Gulf pipefish and stickleback used in this publication. This research was funded by the National Science Foundation Grant OPP-2015301 (to WAC, SB, and Clayton Small), University of Oregon Research Excellence funds (WAC) and National Institute of Health Fellowship 5F31DE032559-02 (to HMH). Additionally, HMH was supported by the Genetics Training Program (NIH T32GM007413).

## Author Contributions

**Hope Healey**: Conceptualization, Methodology, Software, Validation, Formal analysis, Investigation, Data Curation, Writing – Original Draft, Review, and Editing, Visualization, Funding acquisition, **Clara Rehmann**: Methodology, Writing – Review and Editing, **Susan Bassham**: Conceptualization, Writing – Review and Editing, **William Cresko**: Conceptualization, Resources, Writing – Review and Editing, Supervision, Funding acquisition

