## Supplementary information for "Lineage-specific change in craniofacial gene regulatory networks of syngnathid fishes revealed by integrating multiomics across fishes"

### Figures

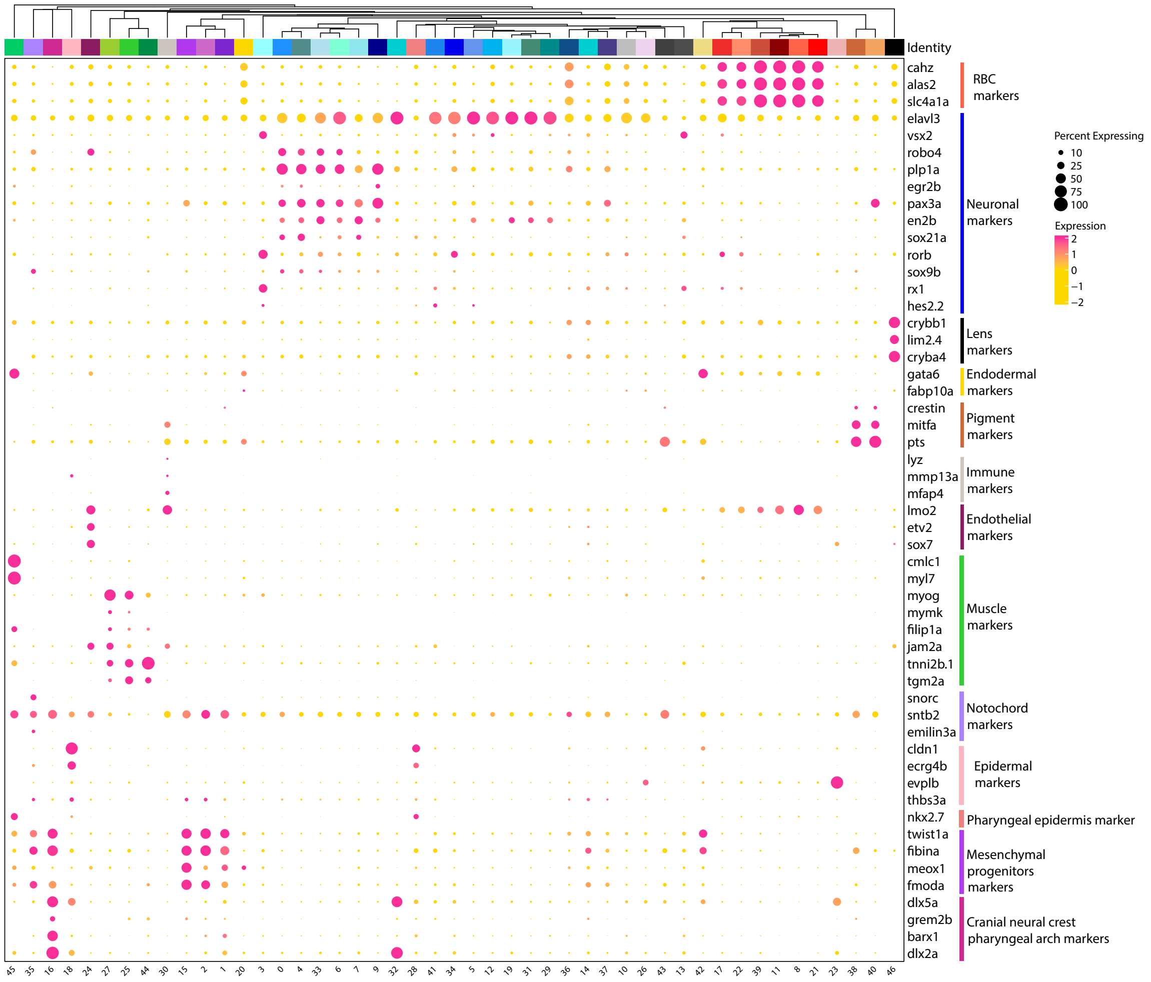


**Fig. S1.** *Marker genes used to define cluster identity*. Using marker genes from the literature and previous zebrafish scRNAseq studies, we inferred cluster identity. Plotted are select markers for each cluster, represented by colored blocks at the top. The size of each dot corresponds to the percentage of cells that express the gene and the dot color denotes the average expression of the gene.

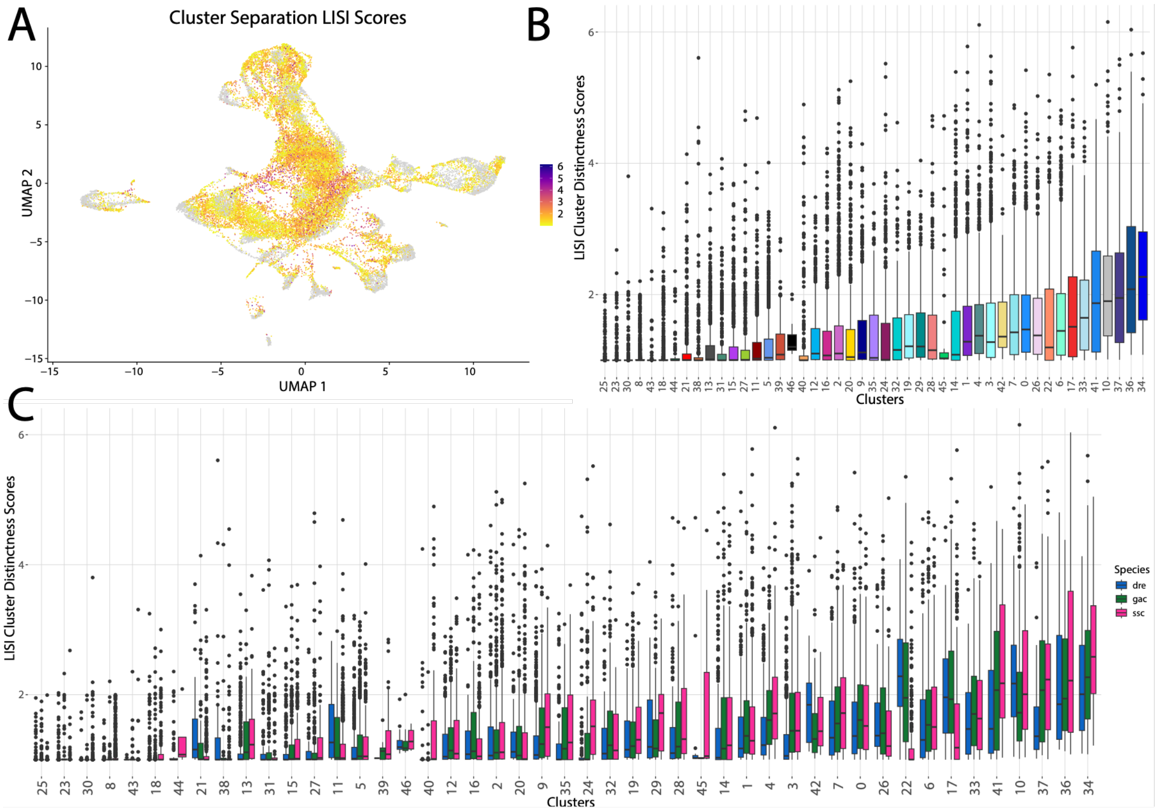


**Fig. S2.** *Comparative scRNAseq atlas reveals a high level of cluster uniqueness*. To evaluate the overlap of cells between clusters, LISI scores were quantified. Scores of 1 indicate cells that are close in UMAP space with only cells of the same cluster. LISI values above 1 represent cells that are nearby cells from other clusters in UMAP space. Scores are relatively similar across the UMAP space (A), with the highest degree of cluster mixing in neural clusters. Cells are plotted in a FeaturePlot in panel A and are colored by the LISI cluster mixing score. Pharyngeal arch cells (#16) form a distinct cell cluster (B) with a low amount of mixing with other clusters. Cluster distinctness minimally varies by species (C). Cluster data are plotted individually in panels B and C, and species data are separated in panel C.

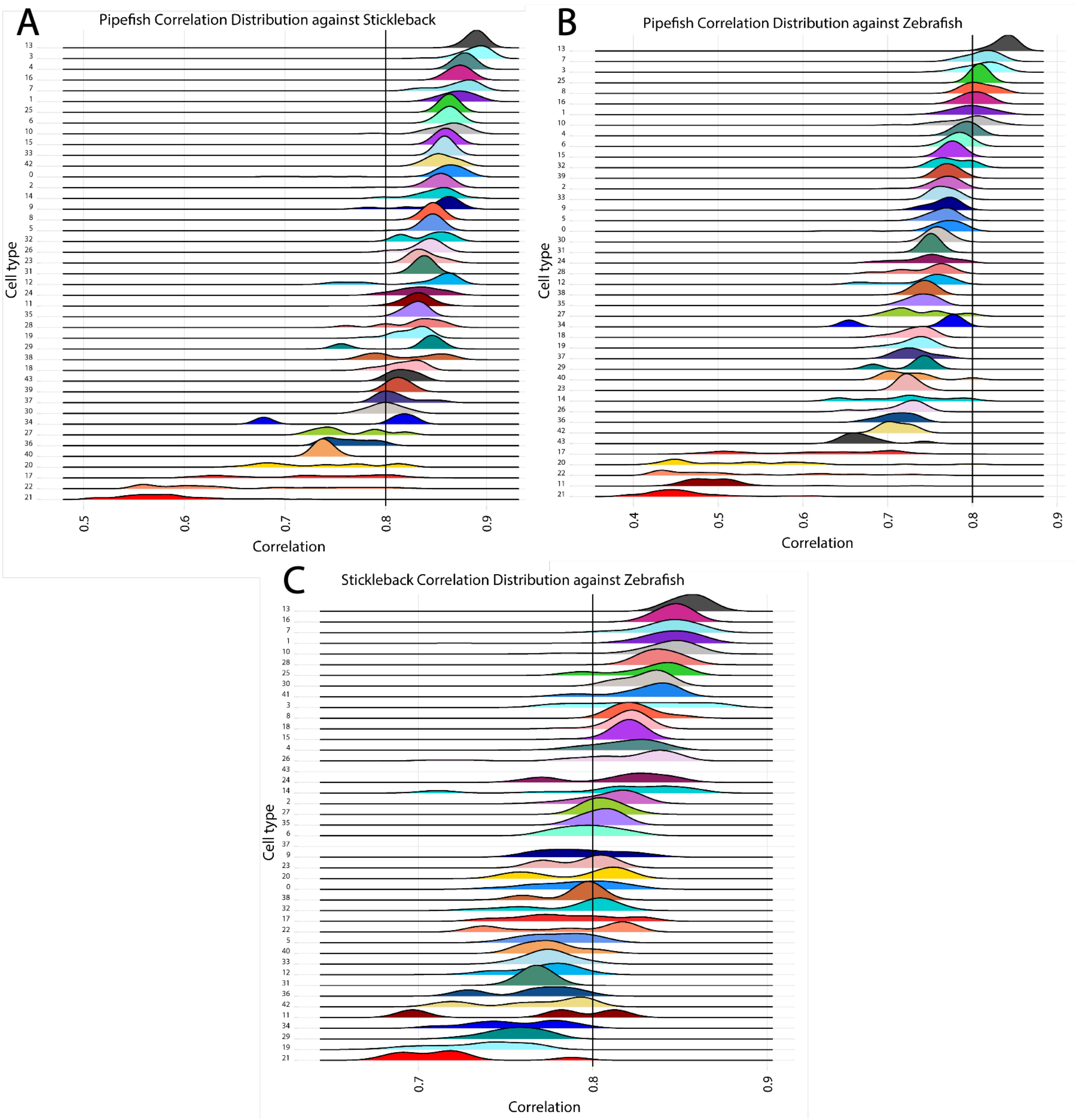


**Fig. S3.** *Pharyngeal arch cells express a highly conserved core genetic program*. In panels A-C, pairwise correlation analyses demonstrate that CNC pharyngeal arch cells (cluster 16) have maintained a core gene expression profile in all species.

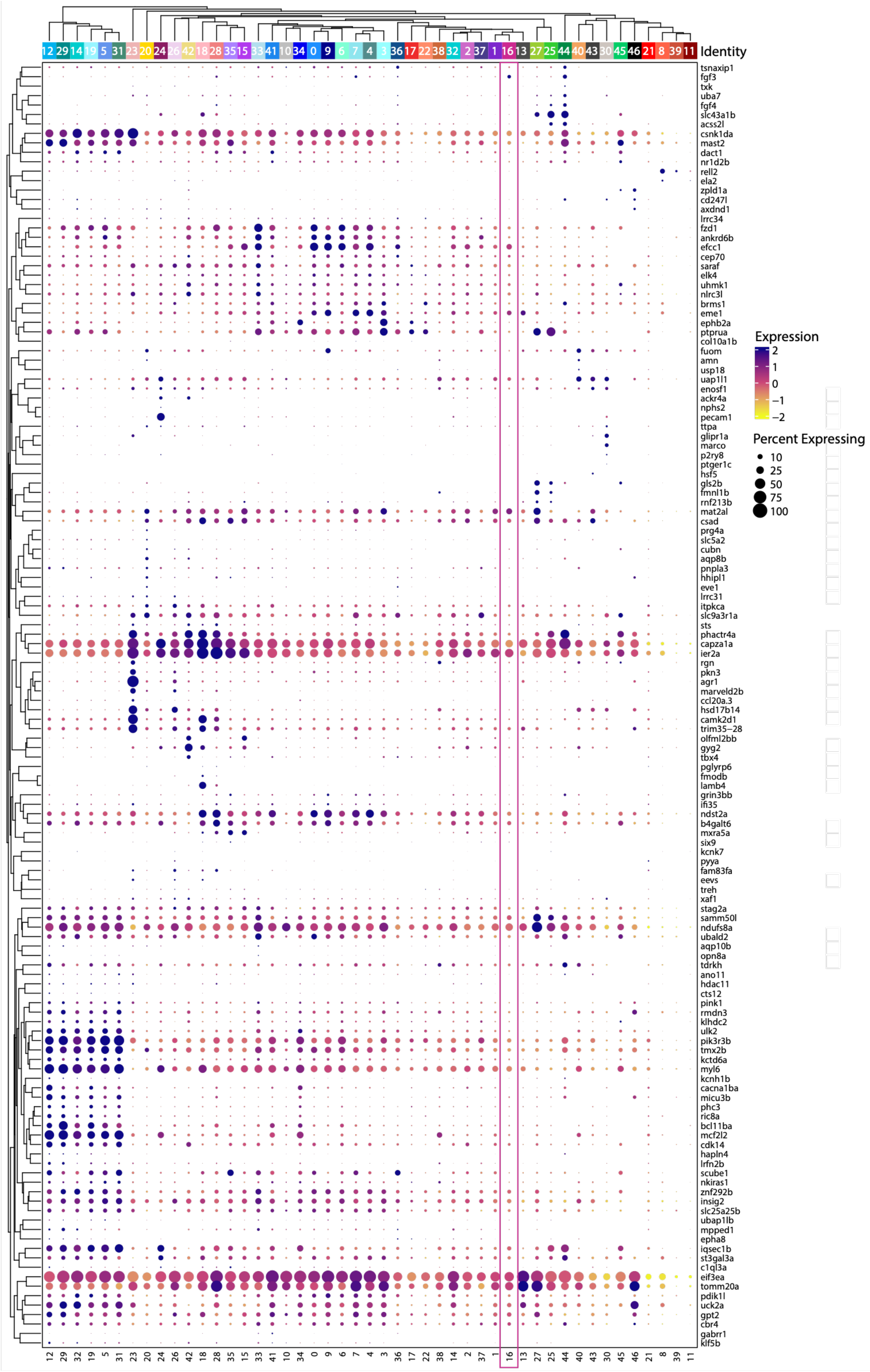


**Fig. S4.** *Expression patterns of genes not identified in syngnathid fishes*. Genes are plotted in a clustered dot plot where the cluster numbers are on the x-axis, gene names are on the y-axis, and both are clustered based on similarity of expression. The size of the dot correlates with the percentage of cells that express it and the hue corresponds to its expression level.

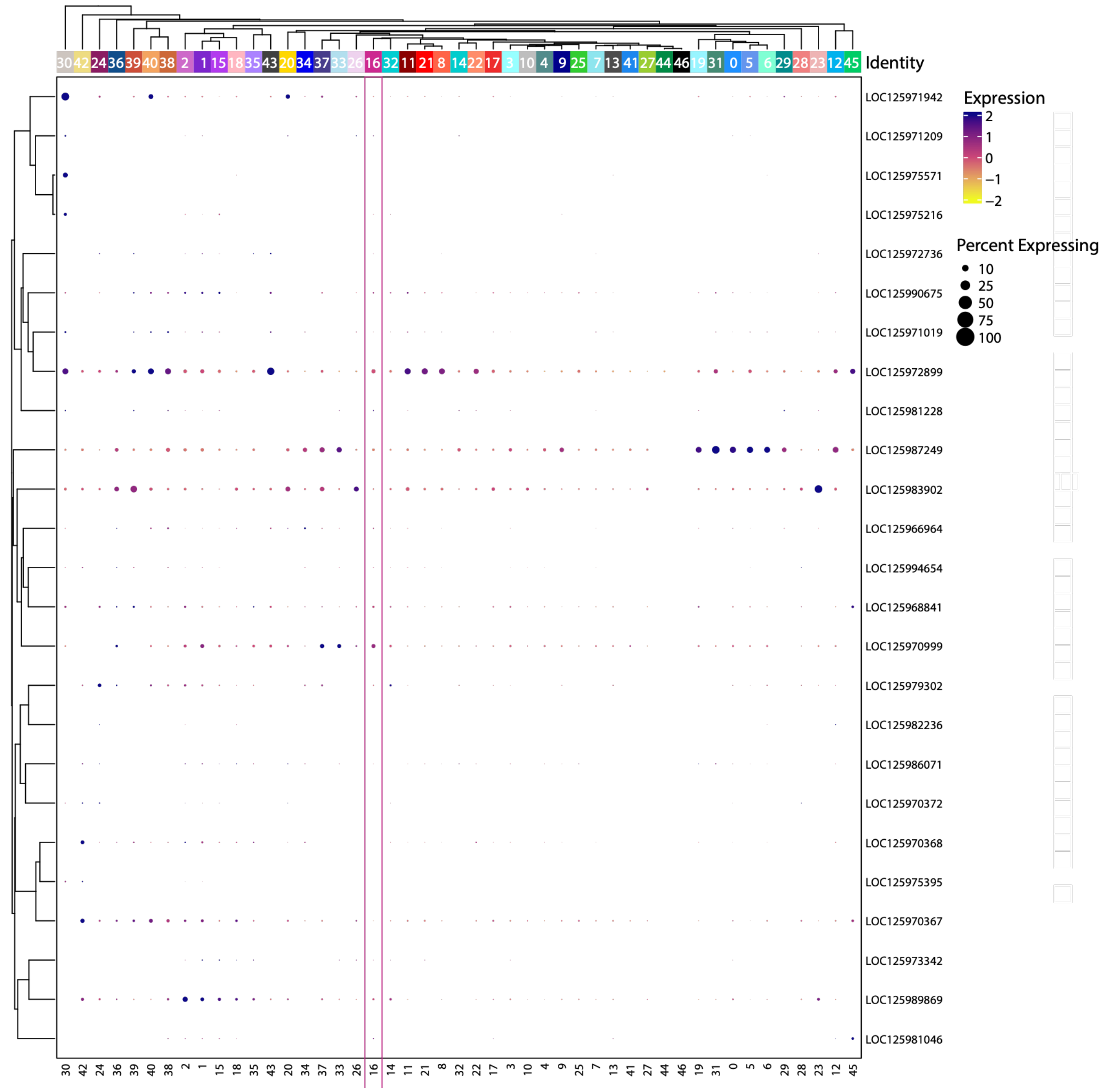


**Fig. S5.** *Expression patterns of genes identified only in sampled syngnathid fishes.* Genes are plotted in a clustered dot plot where the cluster numbers are on the x-axis, gene names are on the y-axis, and both are clustered based on similarity of expression. The size of the dot correlates with the percentage of cells that express it and the hue corresponds to its expression level.

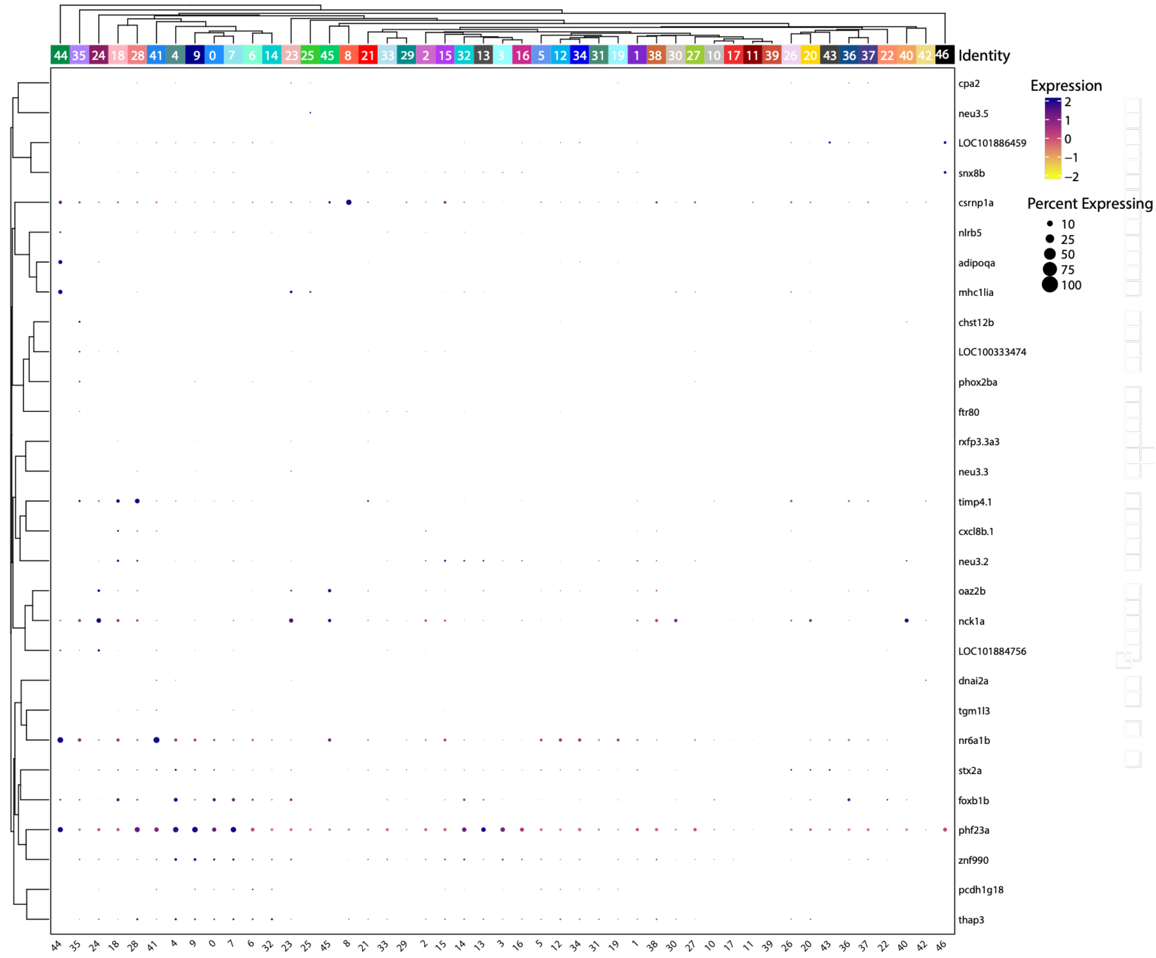


**Fig. S6.** *Expression patterns of genes identified only in sampled cypriniform fishes*. Genes are plotted in a clustered dot plot where the cluster numbers are on the x-axis, gene names are on the y-axis, and both are clustered based on similarity of expression. The size of the dot correlates with the percentage of cells that express it and the hue corresponds to its expression level.

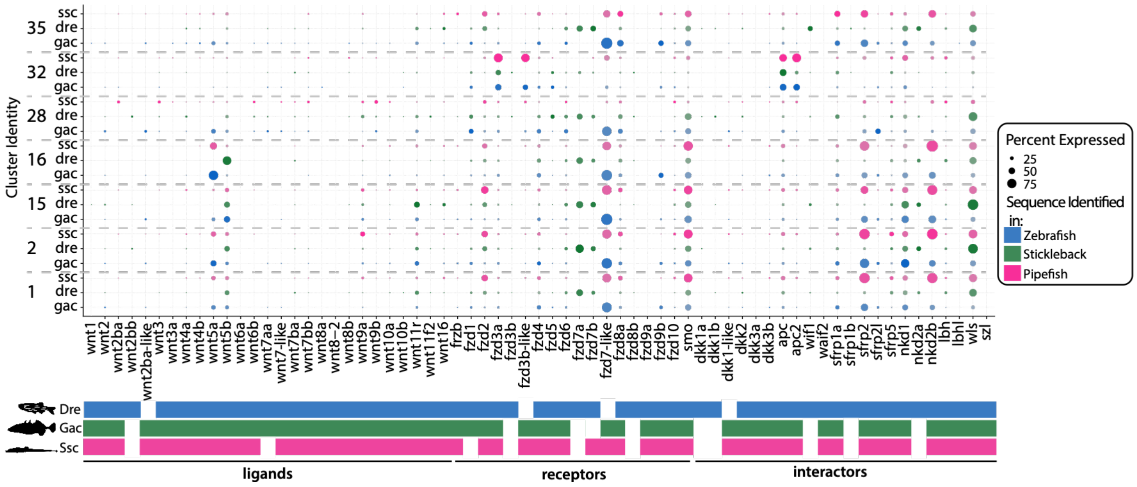


**Fig. S7.** *Wnt pathway in CNC and connective tissue cell types by species*. Clusters are listed on the y-axis and are broken down by species. Wnt ligands, receptors, and interactors are distinguished on the x-axis and are organized by function. Dots represent the percentage of cells for a specific cluster that express each gene. The presence/absence of genes in each species are distinguished in the colored bars below the graph. Empty boxes represent a protein coding gene loss in a species.

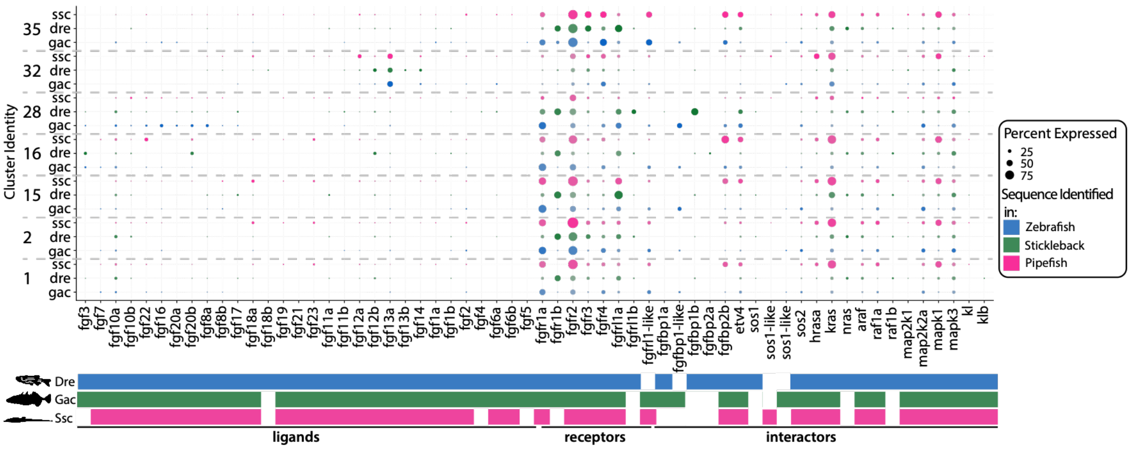


**Fig. S8**. *Fgf pathway in CNC and connective tissue cell types by species.* Clusters are listed on the y-axis and are broken down by species. Fgf ligands, receptors, and interactors are distinguished on the x-axis and are organized by function. Dots represent the percentage of cells for a specific cluster that express each gene. The presence/absence of genes in each species are distinguished in the colored bars below the graph. Empty boxes represent a protein coding gene loss in a species.

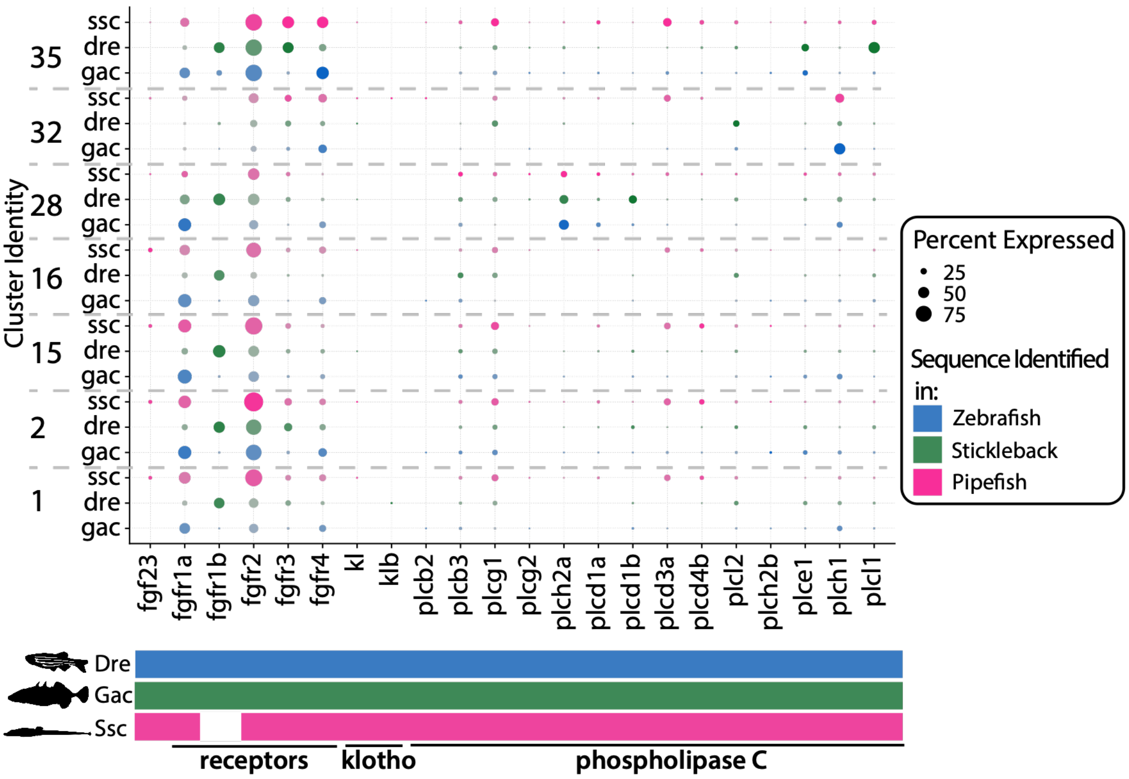


**Fig. S9.** *fgf23, receptors, and potential co-factors in CNC and connective tissue cell types by species*. Clusters are listed on the y-axis and are broken down by species. Fgf receptors and fgf23 co-factors are distinguished on the x-axis and are organized by function. Dots represent the percentage of cells for a specific cluster that express each gene. The presence/absence of genes in each species are distinguished in the colored bars below the graph. Empty boxes represent a protein coding gene loss in a species.

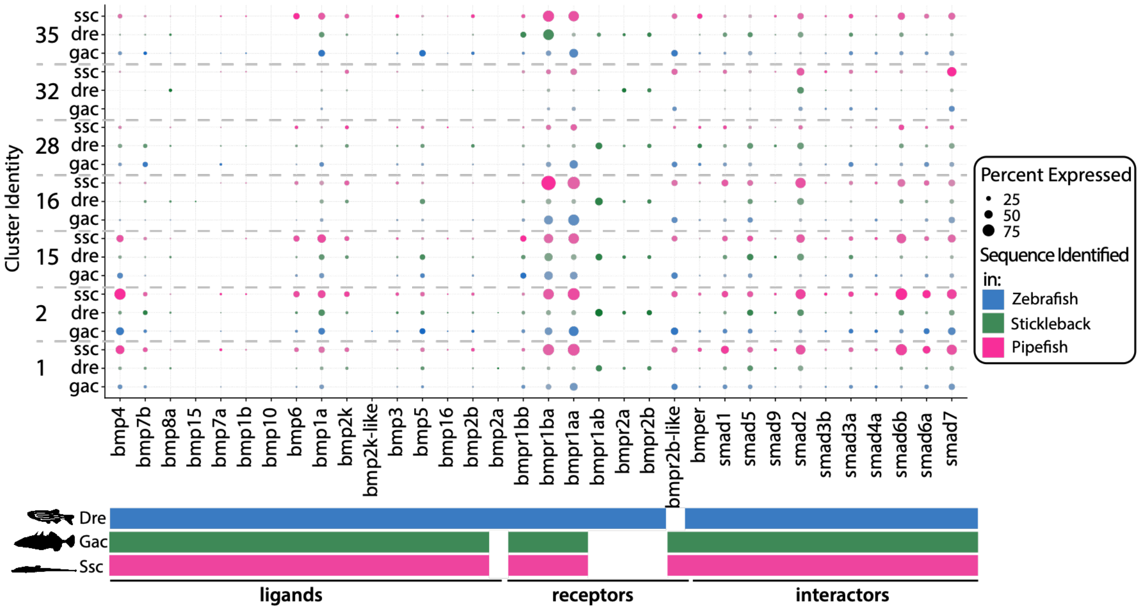


**Fig. S10.** *Bmp pathway in CNC and connective tissue cell types by species*. Clusters are listed on the y-axis and are broken down by species. Bmp ligands, receptors, and interactors are distinguished on the x-axis and are organized by function. Dots represent the percentage of cells for a specific cluster that express each gene. The presence/absence of genes in each species are distinguished in the colored bars below the graph. Empty boxes represent a protein coding gene loss in a species.

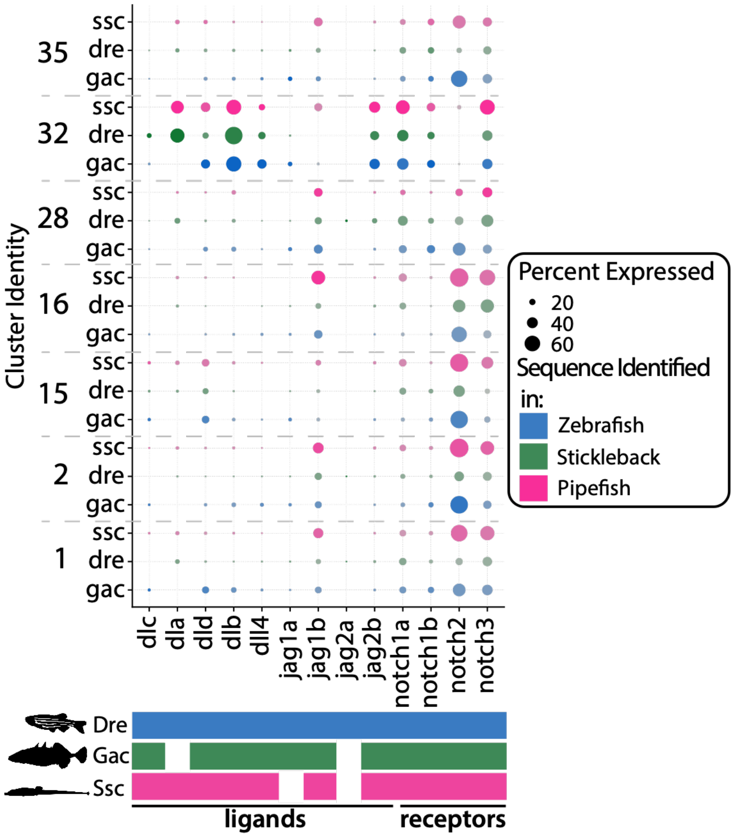


**Fig. S11.** *Notch pathway in CNC and connective tissue cell types by species*. Clusters are listed on the y-axis and are broken down by species. Notch ligands, receptors, and interactors are distinguished on the x-axis and are organized by function. Dots represent the percentage of cells for a specific cluster that express each gene. The presence/absence of genes in each species are distinguished in the colored bars below the graph. Empty boxes represent a protein coding gene loss in a species.

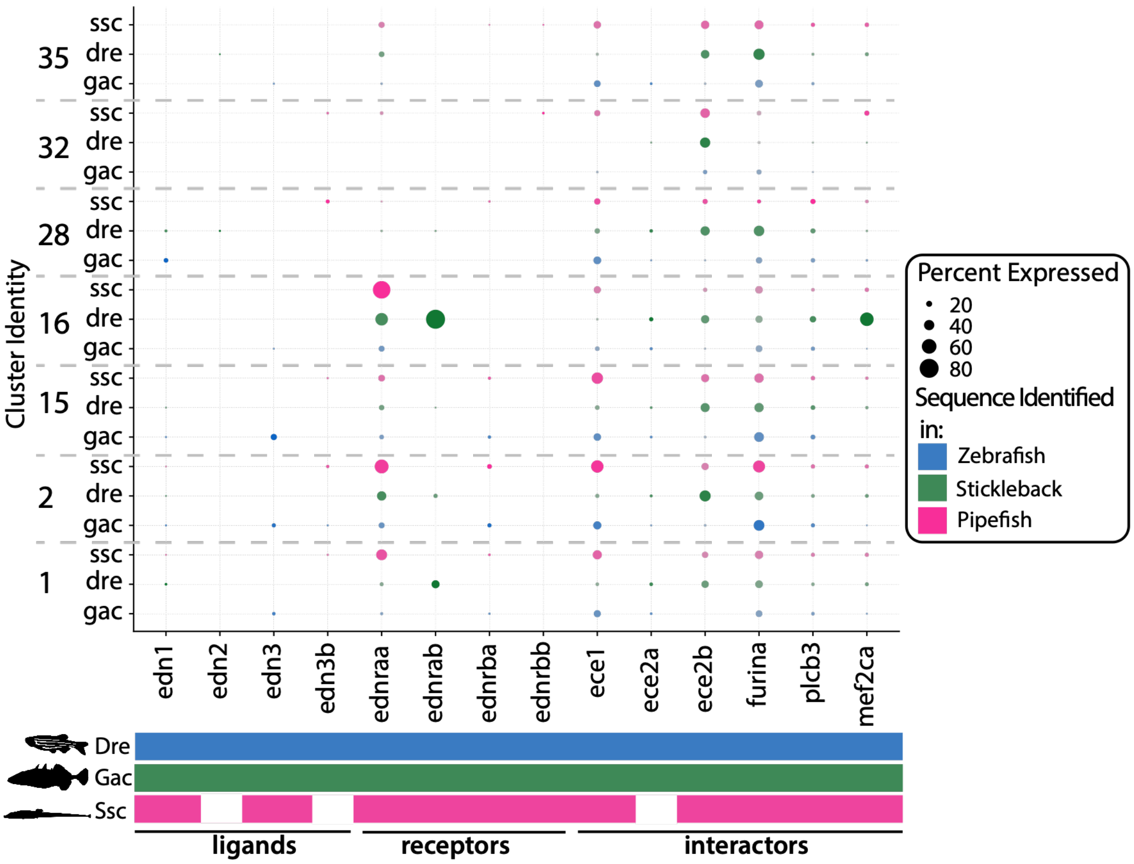


**Fig. S12.** *Endothelin pathway in CNC and connective tissue cell types by species*. Clusters are listed on the y-axis and are broken down by species. Endothelin ligands, receptors, and interactors are distinguished on the x-axis and are organized by function. Dots represent the percentage of cells for a specific cluster that express each gene. The presence/absence of genes in each species are distinguished in the colored bars below the graph. Empty boxes represent a protein coding gene loss in a species.

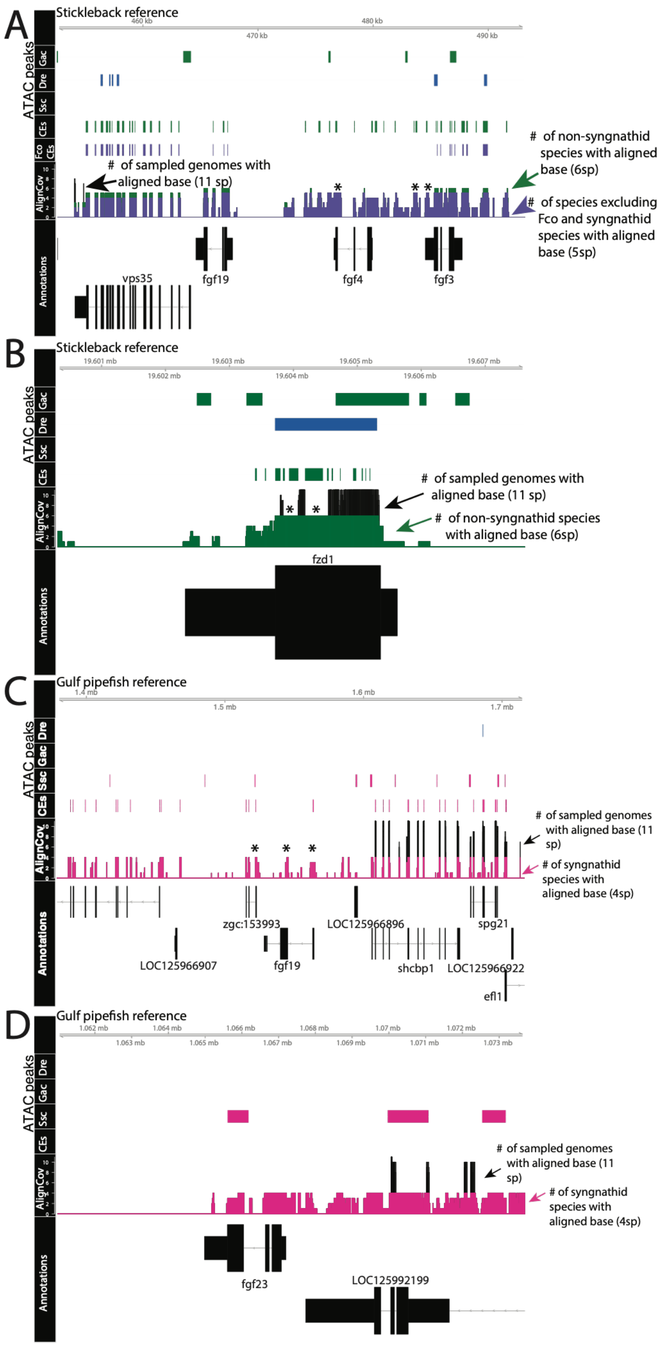


**Fig. S13.** *Changes in conserved elements across teleost evolution.* Panels A-D illustrate identified conserved elements (CE), ATAC-seq peaks, and illustrate genome alignment coverage around signaling genes. Gene models are shown at the base of the graph, whereby thicker regions show exons and thinner regions show introns.

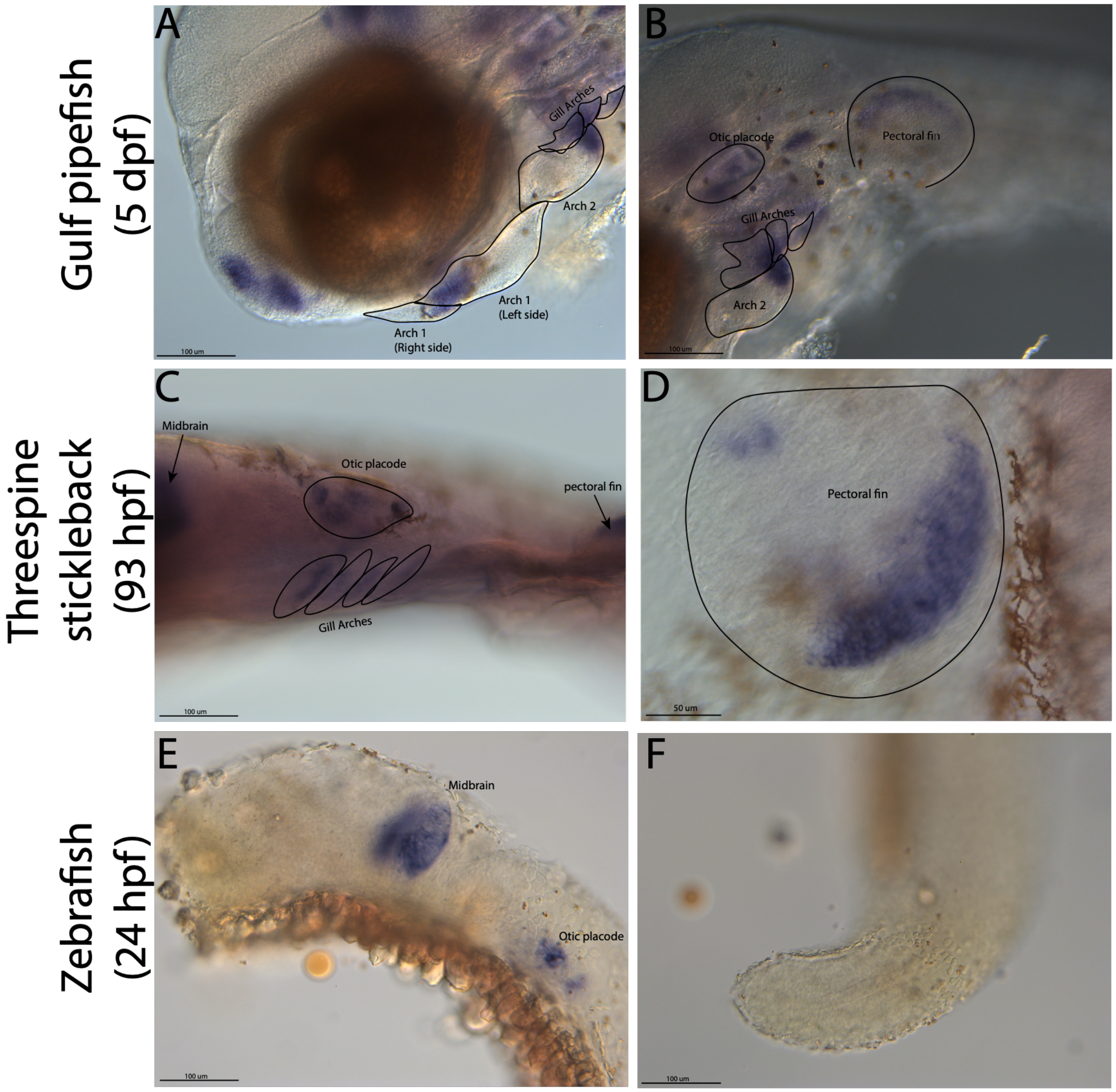


**Fig. S14.** *fgf22 has novel CNC pharyngeal arch and fin expression in percomorphs*. In situ hybridizations from Gulf pipefish, threespine stickleback, and zebrafish reveal spatial gene expression of fgf22. All three fishes had midbrain and otic placode staining (shown in lateral views in A, B, C, E). However, stickleback and pipefish, but not zebrafish, had pharyngeal arch and fin staining (B, D, F). Pharyngeal arch staining of fgf22 was not detected in stickleback via in situ at 70hpf, perhaps due to minimal expression. Fishes were mounted laterally.

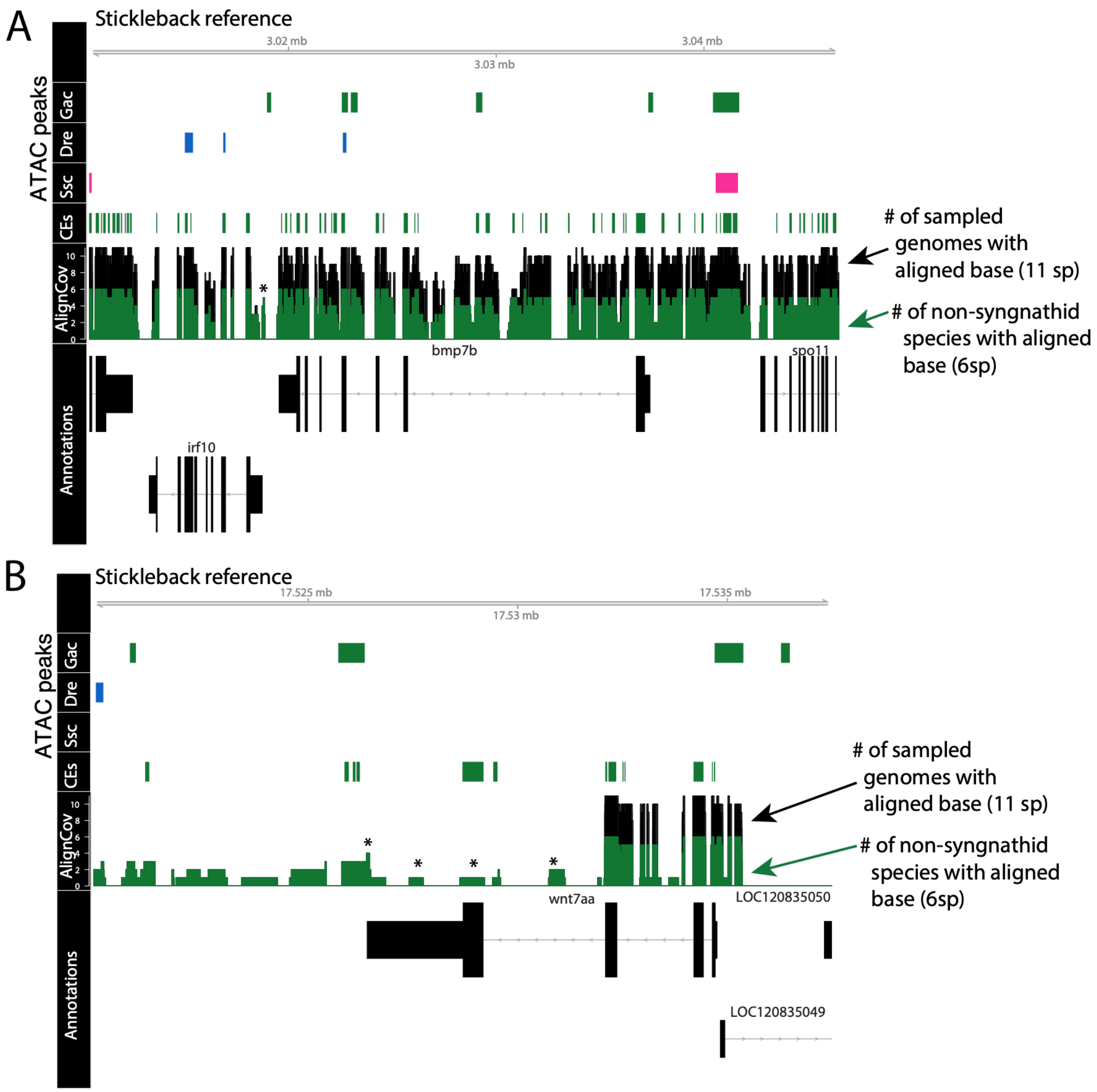


**Fig. S15.** *Potential changes in gene sequence and regulation may lead to Gulf pipefish expression differences.* Panels A and B illustrate identified conserved elements from stickleback (CEs), ATAC-seq peaks, and alignment base pair coverage (align cov). The green plot of alignment coverage shows coverage by base for all species except syngnathids, syngnathids are included in the black plot. Gene models are shown at the base of the graph, whereby thicker regions show exons and thinner regions show introns. Plots illustrate changes in genomic regions in genes that have altered expression or content in Gulf pipefish.

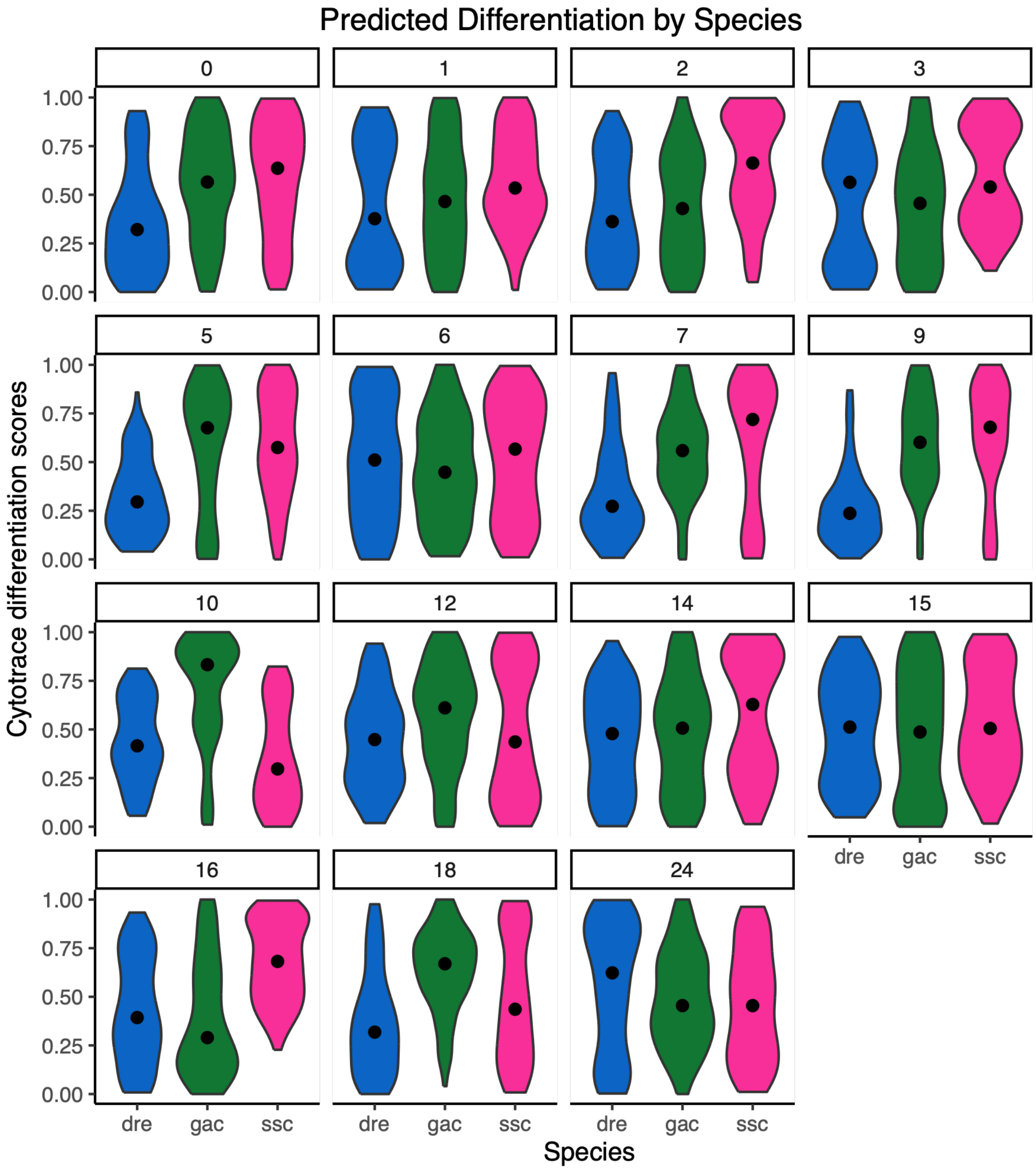


**Fig. S16.** *Species differentiation state varied by cluster though showing no clear evidence for developmental stage mismatch.* Cellular differentiation was tested using integrative CytoTRACE. Species differentiation states for each cell in a cluster are plotted in violin plots. The CytoTRACE scale ranges from 0 to 1, whereby 1 represents undifferentiated cells.

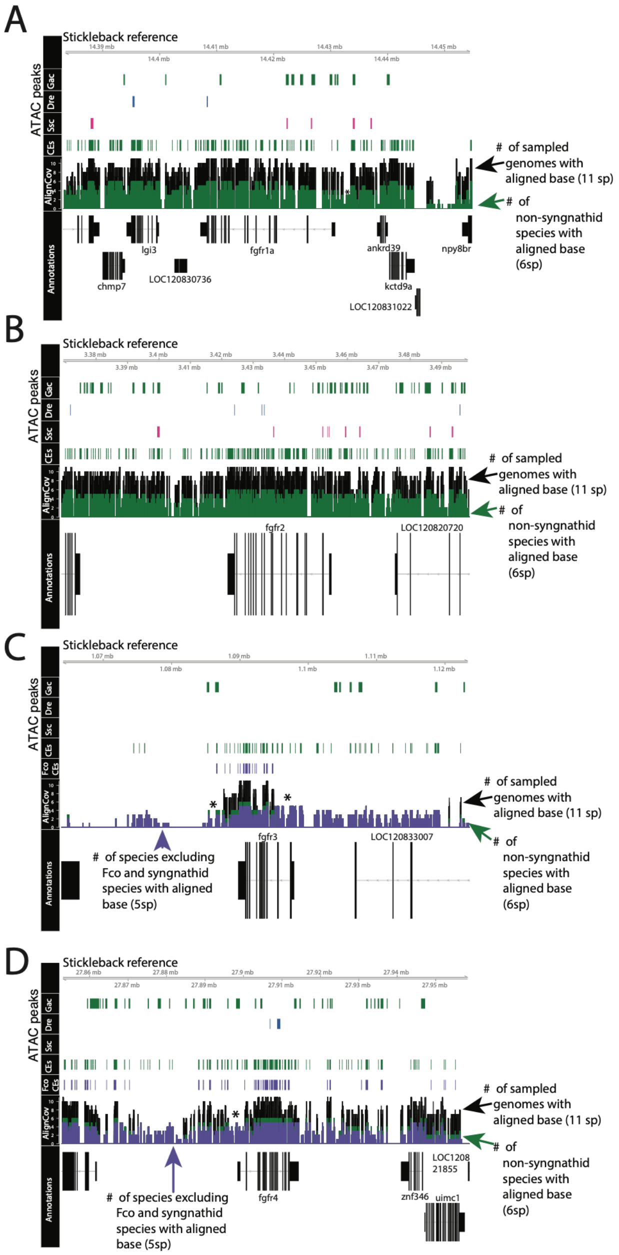


**Fig. S17.** *Potential changes in fgfr gene regulation may lead to Gulf pipefish expression differences*. Panels A-D illustrate identified conserved elements in stickleback (CE), ATAC-seq peaks, and genome alignment coverage. Gene models are shown at the base of the graph, whereby thicker regions show exons and thinner regions show introns. Stickleback is the reference genome for all the plots.

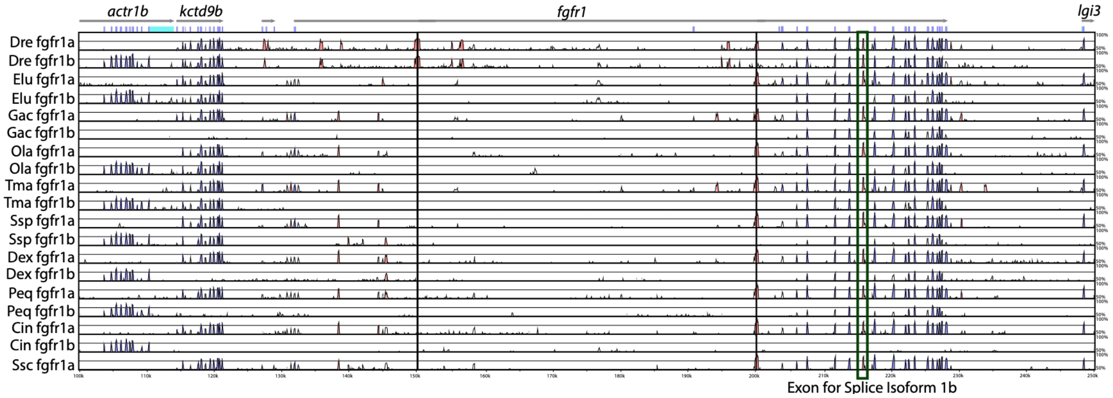


**Fig. S18.** *Splice isoform 1B is missing in northern pike and percomorph fishes fgfr1b*. Species sequence conservation is depicted on a VISTA plot. Exons are shown in dark blue, conserved non-coding elements in orange, and UTRs in light blue. The exon for splice isoform 1b is boxed in cyan. The teleost species are plotted against spotted gar, which has not experienced the teleost whole genome duplication event.

### Tables

**Table S1.** *Correlation medians for species comparisons illustrate high conservation of PA cells*. Median correlation scores from the correlation analysis are listed by species comparison. These assessments are directional and completed for every cluster. The highest score for each species contrast is bolded.

| Cluster | Zebrafish vs Stickleback | Zebrafish vs Pipefish | Stickleback vs Pipefish | Stickleback vs Zebrafish | Pipefish vs Stickleback | Pipefish vs Zebrafish |
|---|---|---|---|---|---|---|
| 0 | 0.81 | 0.78 | 0.86 | 0.79 | 0.86 | 0.77 |
| 1 | 0.85 | 0.82 | 0.88 | 0.85 | 0.87 | 0.80 |
| 2 | 0.81 | 0.78 | 0.86 | 0.81 | 0.85 | 0.77 |
| 3 | **0.86** | 0.81 | 0.88 | 0.83 | **0.89** | 0.81 |
| 4 | 0.83 | 0.79 | 0.88 | 0.82 | 0.88 | 0.79 |
| 5 | 0.78 | 0.76 | 0.84 | 0.78 | 0.85 | 0.77 |
| 6 | 0.81 | 0.78 | 0.86 | 0.80 | 0.86 | 0.79 |
| 7 | 0.85 | 0.81 | 0.88 | 0.84 | 0.88 | 0.81 |
| 8 | 0.84 | 0.82 | 0.85 | 0.82 | 0.85 | 0.81 |
| 9 | 0.82 | 0.77 | 0.86 | 0.79 | 0.86 | 0.77 |
| 10 | **0.86** | 0.82 | 0.86 | 0.85 | 0.87 | 0.80 |
| 11 | 0.81 | 0.73 | 0.86 | 0.78 | 0.83 | 0.49 |
| 12 | 0.78 | 0.75 | 0.85 | 0.78 | 0.86 | 0.75 |
| 13 | **0.86** | **0.84** | **0.89** | **0.86** | **0.89** | **0.84** |
| 14 | 0.84 | 0.79 | 0.86 | 0.82 | 0.85 | 0.72 |
| 15 | 0.82 | 0.77 | 0.86 | 0.82 | 0.86 | 0.78 |
| 16 | 0.85 | 0.82 | 0.88 | 0.85 | 0.87 | 0.80 |
| 17 | 0.83 | 0.77 | 0.80 | 0.78 | 0.74 | 0.63 |
| 18 | 0.82 | 0.74 | 0.83 | 0.82 | 0.82 | 0.74 |
| 19 | 0.77 | 0.74 | 0.83 | 0.74 | 0.83 | 0.74 |
| 20 | 0.78 | 0.60 | 0.77 | 0.81 | 0.74 | 0.52 |
| 21 | 0.75 | 0.59 | 0.63 | 0.71 | 0.58 | 0.45 |
| 22 | 0.81 | 0.76 | 0.82 | 0.79 | 0.62 | 0.49 |
| 23 | 0.76 | 0.69 | 0.84 | 0.80 | 0.84 | 0.72 |
| 24 | 0.79 | 0.75 | 0.83 | 0.82 | 0.84 | 0.75 |
| 25 | 0.83 | 0.80 | 0.85 | 0.84 | 0.86 | 0.81 |
| 26 | 0.81 | 0.71 | 0.83 | 0.83 | 0.84 | 0.73 |
| 27 | 0.81 | 0.78 | 0.71 | 0.80 | 0.75 | 0.72 |
| 28 | 0.84 | 0.76 | 0.84 | 0.84 | 0.84 | 0.75 |
| 29 | 0.76 | 0.73 | 0.82 | 0.76 | 0.84 | 0.74 |
| 30 | 0.84 | 0.78 | 0.83 | 0.84 | 0.80 | 0.76 |
| 31 | 0.77 | 0.75 | 0.84 | 0.77 | 0.84 | 0.75 |
| 32 | 0.80 | 0.76 | 0.83 | 0.80 | 0.85 | 0.77 |
| 33 | 0.79 | 0.77 | 0.85 | 0.77 | 0.86 | 0.76 |
| 34 | 0.78 | 0.77 | 0.78 | 0.75 | 0.81 | 0.77 |
| 35 | 0.80 | 0.73 | 0.84 | 0.81 | 0.83 | 0.74 |
| 36 | 0.78 | 0.68 | 0.75 | 0.77 | 0.76 | 0.71 |
| 37 | 0.78 | 0.75 | 0.85 | 0.79 | 0.81 | 0.73 |
| 38 | 0.78 | 0.74 | 0.87 | 0.80 | 0.82 | 0.74 |
| 40 | 0.78 | 0.74 | 0.74 | 0.78 | 0.74 | 0.72 |

**Table S2.** *Correlation medians for binned species comparisons illustrate high conservation of PA cells*. Median correlation scores from the correlation analysis are listed by species comparison. These assessments are directional and completed for binned clusters. The highest score for each species contrast is bolded.

| | Neural | Conn | RBC | Unk | CNC PA | Epit | Endo | Endot | Musc | Immu | Pigm |
|---|---|---|---|---|---|---|---|---|---|---|---|
| Zebrafish vs Stickleback | 0.83 | 0.83 | 0.84 | **0.86** | 0.85 | 0.82 | 0.78 | 0.79 | 0.83 | 0.84 | 0.78 |
| Zebrafish vs Pipefish | 0.79 | 0.80 | **0.82** | **0.82** | **0.82** | 0.74 | 0.65 | 0.75 | 0.79 | 0.78 | 0.74 |
| Stickleback vs Pipefish | 0.86 | 0.87 | 0.81 | 0.86 | **0.88** | 0.84 | 0.81 | 0.83 | 0.75 | 0.83 | 0.76 |
| Stickleback vs Zebrafish | 0.79 | 0.82 | 0.80 | **0.85** | **0.85** | 0.82 | 0.76 | 0.82 | 0.81 | 0.84 | 0.80 |
| Pipefish vs Zebrafish | 0.78 | 0.78 | 0.80 | 0.80 | 0.80 | 0.73 | 0.54 | 0.75 | **0.81** | 0.76 | 0.74 |
| Pipefish vs Stickleback | 0.86 | 0.86 | 0.84 | **0.87** | **0.87** | 0.83 | 0.77 | 0.84 | 0.86 | 0.80 | 0.79 |

**Table S3.** Orthology renaming scheme.

| ORTHOLOGY (ZEBRAFISH:STICKLEBACK:PIPEFISH) | ZEBRAFISH GENES | STICKLEBACK GENES | PIPEFISH GENES |
|---|---|---|---|
| **1:1:1** | Name based on zebrafish | Name based on zebrafish | Name based on zebrafish |
| **1:0:1** | Name based on zebrafish | NA | Name based on zebrafish |
| **0:1:1** | NA | Name based on stickleback | Name based on stickleback |
| **1:0:0** | Name based on zebrafish | NA | NA |
| **0:1:0** | NA | Name based on stickleback | NA |
| **0:0:1** | NA | NA | Name based on pipefish |

**Dataset S1.** Genomes used in GENESPACE and Cactus analyses.

**Dataset S2.** Gene orthologs identified from the 28 species GENESPACE analysis for zebrafish, stickleback, and pipefish used to rename scRNAseq analysis.

**Dataset S3.** Identified orthologs from GENESPACE with zebrafish (Danio rerio) as the reference species used for gene content analysis

**Dataset S4.** Identified orthologs from GENESPACE with Gulf pipefish (Syngnathus scovelli) as the reference species used for gene content analysis

**Dataset S5.** Identified orthologs from Genespace with stickleback (Gasterosteus aculeatus) as the reference species used for gene content analysis

**Dataset S6.** Cluster annotations for the integrated atlas.

**Dataset S7.** Genes that were not identified in any syngnathids from the GENESPACE analysis that were present in a close outgroup, mandarin dragonet, and the two other focal species (zebrafish and stickleback).

**Dataset S8.** Genes that were not identified in any perciformes from the GENESPACE analysis that were present in a close outgroup, fugu, and the two other focal species (zebrafish and Gulf pipefish).

**Dataset S9.** Genes that were not identified in any cypriniform from the GENESPACE analysis that were present in a close outgroup, Mexican blind cavefish, and the two other focal species (stickleback and Gulf pipefish).

**Dataset S10.** Genes that were only detected in syngnathids.

**Dataset S11.** Genes that were only detected in cypriniformes.

**Dataset S12.** File containing all syngnathid candidate genes identified through CE, scRNAseq, and GENESPACE analysis.

**Dataset S13.** Cytotrace analysis ran to test developmental stage of cells from different species.
